# A dual-inducible control system for multistep biosynthetic pathways

**DOI:** 10.1101/2024.06.12.598700

**Authors:** Andrés Felipe Carrillo Rincón, Alexandra J. Cabral, Andras Gyorgy, Natalie G. Farny

## Abstract

**Background:** The successful production of industrially relevant natural products hinges on two key factors: the cultivation of robust microbial chassis capable of synthesizing the desired compounds, and the availability of reliable genetic tools for expressing target genes. The development of versatile and portable genetic tools offers a streamlined pathway to efficiently produce a variety of compounds in well-established chassis organisms. The σ^70^ *lac* and *tet* expression systems – adaptations of the widely used *lac* and *tet* regulatory systems developed in our laboratory – have shown effective regulation and robust expression of recombinant proteins in various Gram-negative bacteria. Understanding the strengths and limitations of these regulatory systems in controlling recombinant protein production is essential for progress in this area.

**Results:** To assess their capacity for combinatorial control, both the σ^70^ *lac* and *tet* expression systems were combined into a single plasmid and assessed for their performance in producing fluorescent reporters as well as the terpenoids lycopene and β-carotene. We thoroughly characterized the induction range, potential for synergistic effects, and metabolic costs of our dual σ^70^ *lac* and *tet* expression system in the well-established microorganisms *Escherichia coli*, *Pseudomonas putida*, and *Vibrio natriegens* using combinations of fluorescent reporters. The dynamic range and basal transcriptional control of the σ^70^ expression systems were further improved through the incorporation of translational control mechanisms via toehold switches. This improvement was assessed using the highly sensitive luciferase reporter system. The improvement in control afforded by the integration of the toehold switches enabled the accumulation of a biosynthetic intermediate (lycopene) in the β-carotene synthesis pathway.

**Conclusion:** This study presents the development and remaining challenges of a set of versatile genetic tools that are portable across well-established gammaproteobacterial chassis and capable of controlling the expression of multigene biosynthetic pathways. The enhanced σ^70^ expression systems, combined with toehold switches, facilitate the biosynthesis and study of enzymes, recombinant proteins, and natural products, thus providing a valuable resource for producing a variety of compounds in microbial cell factories.

**GRAPHICAL ABSTRACT:** 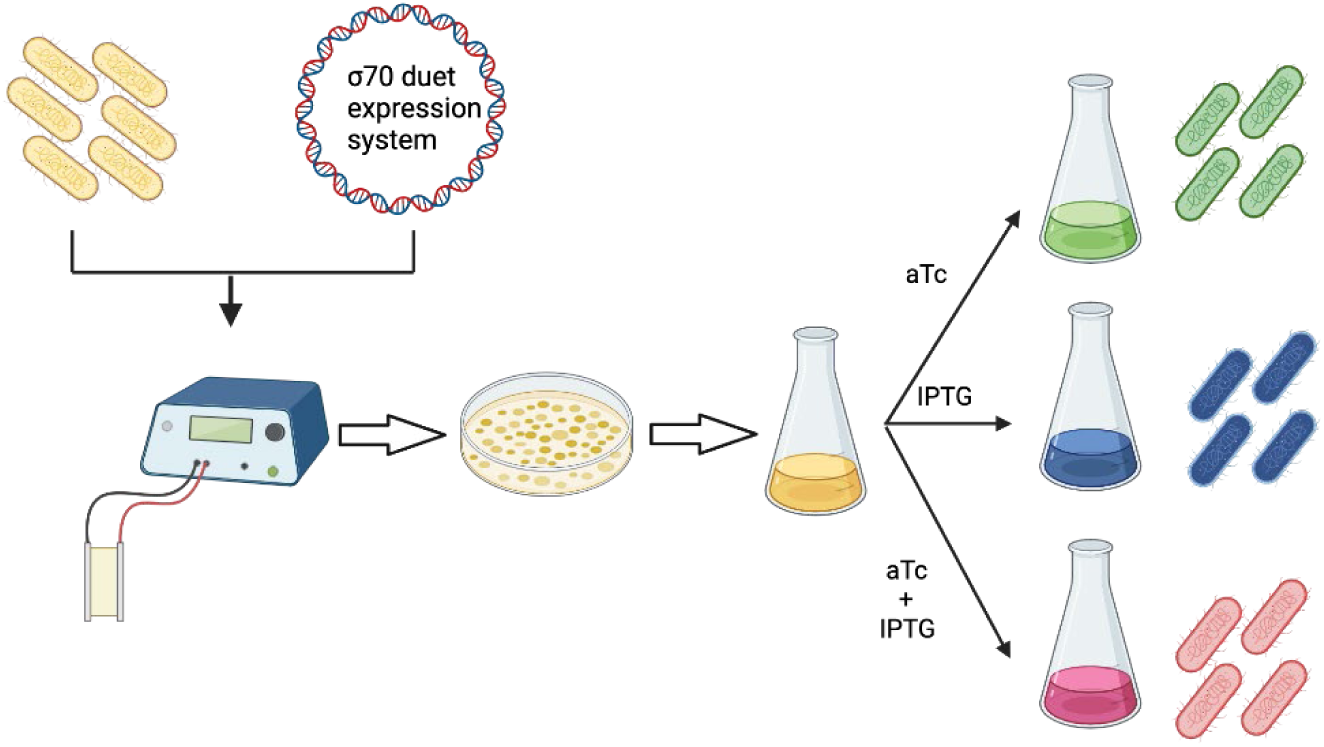

## BACKGROUND

Modern sequencing techniques have unveiled a myriad of unknown compounds and proteins encoded by microorganisms with potential industrial applications ranging from biotherapeutics to sustainable manufacturing [1, 2]. However, these microorganisms are not always easy to cultivate, and manipulating their genes poses a central challenge in unlocking their full potential [3]. *Escherichia coli* is typically the primary choice for expressing proteins from other organisms due to its robust genetic toolkit and well-established cultivation parameters [4]. In some cases, *E. coli* may struggle to produce toxic genes, membrane proteins, or other complex proteins necessary for the synthesis of natural products [5, 6]. To overcome these limitations of *E. coli*, alternative hosts like *Pseudomonas putida* and *Vibrio natriegens* have become appealing chassis for producing recombinant proteins. They offer standard laboratory techniques akin to *E. coli*, along with robust genetic and metabolic backgrounds, and rapid growth characteristics [7, 8]. Importantly, these microorganisms have successfully produced natural products such as non-ribosomal peptides, polyketides, terpenoids, fatty acids, and polymers [9–13], and can be genetically engineered with a diverse set of genetic tools [14, 15].

The *lac* and *tet* inducible promoters have been workhorses of molecular biology for decades [16–19]. Many studies have been completed to optimize their performance [20, 21] and expand their functionality beyond *E. coli* [14, 22]. Our recent work contributed to this body of research by optimizing the inducible *tet* and *lac* promoters for σ^70^ recognition, showing precise regulation and robust gene overexpression in *E. coli*, *P. putida* and *V. natriegens* [23]. However, we have only assessed our genetic tools for their efficacy in producing the reporter protein *mCardinal* and the cocaine esterase *CocE* [23]. Multiple biological processes necessitate the coordinated activity of numerous genes, such as those involved in synthesizing natural products encoded by biosynthetic gene clusters (BGCs) [24, 25]. To facilitate multi-gene expression, the pET system was modified to concurrently express various genes within the same *E. coli* BL21 strain using the pETDuet system [26]. However, in pETDuet, both multiple cloning sites are induced by IPTG, preventing independent or sequential control [27]. To overcome this limitation, a dual inducible plasmid was developed to independently control gene expression with the *lac* and *P_BAD_* promoters [28]. In addition to the single-plasmid dual-expression system, the *E. coli* “marionette” strain was developed with twelve inducible systems [25]. The “marionette” strain contains the transcriptional regulators integrated in the chromosome, including *lacI*, *tetR* and *araC*, and the target gene is deployed in a replicative plasmid under the control of the respective inducible promoter [25]. Further, the synergistic effect of combined promoters to produce a single target recombinant protein in *E. coli* has demonstrated differing levels of success thus far; while positive synergy has been observed in the production of specific recombinant proteins, the opposite trend has also been reported [29]. A dual-inducible duet expression system was also engineered in *P. putida* using a multicopy plasmid and utilizing the original configuration of the *lac*, *tet* and *P_BAD_* promoters, resulting in the successful production of fluorescent proteins and an inducible CRISPR interference system [30, 31]. Despite these significant achievements, the native promoters utilized in these studies lack the strength and control of the optimized σ^70^ *lac* and *tet* promoter, which we demonstrated enable strong production of recombinant proteins upon induction while maintaining low basal levels in the uninduced state in *E. coli*, *P. putida* and *V. natriegens* [23]. We believe our tools could be useful in improving control and production titer in industrial biotechnology applications, however their practical utility, strengths, and limitations for such applications has not yet been measured.

In our prior work, adapting the consensus sequences recognized by the σ^70^ factor to the *lac* and *tet* promoters significantly enhanced their strength, surpassing the efficiency of expression from the pET system in *E. coli* [23]. Nevertheless, the uninduced state of the optimal σ^70^V2LacI expression system was scarcely repressed by its transcriptional regulator *LacI*, thereby negating the advantages of having an inducible system [23]. One strategy to enhance the fold-change and minimize the leaky expression of the target gene, thereby reducing unnecessary consumption of cellular resources required for cell growth, involves the adaptation of toehold switch riboregulators [32]. In the toehold-mediated strand displacement (TMSD) strategy, the toehold sequesters the ribosome binding site (RBS) of an mRNA molecule through the formation of a stem loop, and translation is only activated in the presence of the trigger RNA complementary to the toehold sequence that linearizes the stem loop [33]. Toehold switches have been shown to significantly enhance the mean dynamic range of reporter expression by over 400-fold [34]. Thus, incorporating toehold switches into strong inducible promoters, like the σ^70^V2LacI, could effectively reduce leakage in the uninduced state, improving the usefulness of this promoter in controlling recombinant protein expression.

In this study, we showed that the strong inducible promoters σ^70^V2Tc and σ^70^V2/3Lac, recognized by the σ^70^, can be adapted into a dual-inducible duet expression system, facilitating the overexpression of multiple genes in well-studied microorganisms such as *E. coli*, *P. putida*, and *V. natriegens*. The efficacy of this duet expression system was confirmed through the expression of the reporter systems *mCardinal* and *sfGFP*, as well as the production of terpenoids lycopene and β-carotene. Furthermore, we enhanced the dynamic range and minimized the leakiness of these expression systems by incorporating toehold switches, a refinement validated by the sensitive multigene reporter *lux* operon (*luxCDABE*). Our toehold-controlled dual inducible system permitted partial accumulation of lycopene as a pathway intermediate in the biosynthesis of β-carotene, which was fully converted to β-carotene with the second inducer, providing proof-of-concept that such control is possible. In summary, our findings illustrate that the σ^70^ duet *lac/tet* expression system offers a versatile gene tool for activating multiple genes with various inducers across different bacterial chassis. Additionally, the incorporation of a toehold switch effectively reduces the expression of target genes in the uninduced state, a crucial feature for sensitive or toxic applications.

## METHODS

### Bacterial strains and culture conditions

*E. coli* DH10B, *P. putida* JE90 [35] and *V. natriegens* [8] were cultured in LB media supplemented with Kanamycin when required (50 µg/mL for *E. coli* and *P. putida*, and 200 µg/mL for *V. natriegens*). *E. coli* and *P. putida* cells were transformed by electroporation, and *V. natriegens* by chemical transformation with the plasmids listed in Table 1. *E. coli* was cultivated at 37°C and *P. putida* and *V. natriegens* at 30°C at 220 rpm.

### Plasmid construction

Phusion® High Fidelity Polymerase (Thermo Scientific) and primers synthesized by Integrated DNA Technologies (IDT) were used in all PCR amplifications for plasmid construction. Plasmids were constructed using Gibson Assembly EX Cloning Kit or T4 DNA ligase (Thermo Scientific) according to manufacturer’s instructions. Plasmids were transformed into ElectroMAX DH10B Cells (Thermo Scientific) according to manufacturer’s instructions. Transformants were selected on LB (Miller) agar plates containing 50 mg/L kanamycin sulfate for selection and incubated at 37°C. Plasmids were constructed using a combination of the ligation of phosphorylated oligonucleotides, and Gibson Assembly. Sequences of all plasmids were confirmed using Sanger sequencing performed by Quintarabio, and/or whole plasmid sequencing by Plasmidsaurus. The details of the molecular cloning steps are available in the supplementary information. The complete list of plasmids used and generated in this study is presented in Table 1. Primers and DNA sequences used in this study are presented in the supplementary information.

**Table 1.** List of plasmids evaluated in this study.

| Name | Expression system<br>1 | ORF1 | Expression system<br>2 | ORF2 | Source |
| --- | --- | --- | --- | --- | --- |
| p$\sigma^{70}$ V2TcR-mCardinal | V2TcR | *mCardinal* | | | [23] |
| p$\sigma^{70}$ V2lacI-mCardinal | V2lacI | *mCardinal* | | | [23] |
| p$\sigma^{70}$ V3LacI-mCardinal | V3LacI | *mCardinal* | | | [23] |
| p$\sigma^{70}$ V2TcR-sfGFP | V2TcR | *sfGFP* | | | [23] |
| p$\sigma^{70}$ V2lacI-sfGFP | V2LacI | *sfGFP* | | | [23] |
| p$\sigma^{70}$ V3LacI-sfGFP | V3LacI | *sfGFP* | | | [23] |
| pJH0204F |  |  |  |  | This study |
| p$\sigma^{70}$ V2TcR-mCardinal/V2LacI-mCardinal | V2TcR | *mCardinal* | V2LacI | *mCardinal* | This study |
| p$\sigma^{70}$ V2TcR-mCardinal/V3LacI-mCardinal | V2TcR | *mCardinal* | V3LacI | *mCardinal* | This study |
| p$\sigma^{70}$ V2TcR- sfGFP/V2LacI-mCardinal | V2TcR | *mCardinal* | V2LacI | *sfGFP* | This study |
| p$\sigma^{70}$ V2TcR- sfGFP/V3LacI-mCardinal | V2TcR | *mCardinal* | V3LacI | *sfGFP* | This study |
| pUC19-crtEBI | - | *crtEBI* |  |  | This study |
| p$\sigma^{70}$ V2TcR-crtEBI | V2TcR | *crtEBI* | | | This study |
| p$\sigma^{70}$ V2TcR-crtEBIY | V2TcR | *crtEBIY* | | | This study |
| p$\sigma^{70}$ V2TcR-crtEBI/V2LacI-crtY | V2TcR | *crtEBI* | V2LacI | *crtY* | This study |
| p$\sigma^{70}$ V2TcR-crtEBI/V3LacI-crtY | V2TcR | *crtEBI* | V3LacI | *crtY* | This study |
| p$\sigma^{70}$ V2TcR-THS-mCardinal | V2TcR | mCardinal | | | This study |
| p$\sigma^{70}$ V3-2LacI-THS-mCardinal | V3-2LacI | mCardinal | | | This study |
| p$\sigma^{70}$ V2TcR-THS-luxCDABE | V2TcR | *luxCDABE* | | | This study |
| p$\sigma^{70}$ V3-2LacI-THS-luxCDABE | V3-2LacI | *luxCDABE* | | | This study |
| p$\sigma^{70}$ V2TcR -luxCDABE | V2TcR | *luxCDABE* | | | This study |
| p$\sigma^{70}$ V2LacI-luxCDABE | V2LacI | *luxCDABE* | | | This study |
| pSC101-Kn |  |  |  |  | This study |
| pSC101Kn $\sigma^{70}$ V2TcR -luxCDABE | V2TcR | *luxCDABE* | | | This study |
| pSC101Kn $\sigma^{70}$ V2LacI-luxCDABE | V2LacI | *luxCDABE* | | | This study |
| pSC101Kn $\sigma^{70}$ V2TcR-THS-luxCDABE | V2TcR | *luxCDABE* | | | This study |
| pSC101Kn $\sigma^{70}$ V3-2LacI-THS-luxCDABE | V3-2LacI | *luxCDABE* | | | This study |
| pSC101Kn $\sigma^{70}$ V2TcR-THS-crtEBI | V2TcR | *crtEBI* | | | This study |
| pSC101Kn $\sigma^{70}$ V2TcR-THS-crtEBIY | V2TcR | *crtEBIY* | | | This study |
| pSC101Kn $\sigma^{70}$ V3-2LacI-THS-crtEBI | V3-2LacI | *crtEBI* | | | This study |
| pSC101Kn $\sigma^{70}$ V3-2LacI-THS-crtEBIY | V3-2LacI | *crtEBIY* | | | This study |
| p$\sigma^{70}$ V2TcR-THS-crtEBI | V2TcR | *crtEBI* | | | This study |
| p$\sigma^{70}$ V2TcR-THS-crtEBIY | V2TcR | *crtEBIY* | | | This study |
| p$\sigma^{70}$ V3-2LacI-THS-crtEBI | V3-2LacI | *crtEBI* | | | This study |
| p$\sigma^{70}$ V3-2LacI-THS-crtEBIY | V3-2LacI | *crtEBIY* | | | This study |
| pSC101Kn $\sigma^{70}$ V2TcR-THS-crtY V3-2LacI-THS-crtEBI | V2TcR | *crtY* | V3-2LacI | *crtEBI* | This study |

### Measurement of the reporter systems in the plate reader

Growth curves and fluorescence measurements were performed in 96-well clear bottom microplates (Corning Incorporated, NY) using the BioTek Synergy H1 Hybrid Multi-Mode Reader following the experimental approach described in [36]. Briefly, overnight cultures of the recombinant strains were diluted into fresh LB medium supplemented with kanamycin to reach OD_600_ 0.05. *E. coli* experiments were performed at 37°C, and *P. putida* and *V. natriegens* at 30°C, shaking at 355 cycles per minute (cpm). Cell density (OD_600_) and fluorescence (*sfGFP* excitation 485nm, emission 510nm; *mCardinal* excitation 604nm, emission 659nm) were monitored every hour for 16 hours. *E. coli* cultures were induced at 3 hours with 0.2 mM IPTG and/or 0.1 µg/mL aTc, *P. putida* at 3 hours with 5 mM IPTG and/or 1 µg/mL aTc, and *V. natriegens* at 2 hours with 0.2 mM IPTG and/or 0.1 µg/mL aTc when required.

### Lycopene and β-carotene determination

Lycopene and β-carotene were measured by UHPLC following the procedure described in [37] using a Shimadzu Nexera UHPLC. Lycopene and β-carotene were extracted with acetone as described in [38] with the following modifications. 5% of overnight cultures of the recombinant *E. coli*, *P. putida* and *V. natriegens* strains containing either the LYC operon or the *crtEBIY* genes were inoculated in 50 mL culture tubes in 5 mL of fresh LB media supplemented with kanamycin and grown at 220 rpm and 37°C for *E. coli* and 30°C for *P. putida* and *V. natriegens*. The cultures were induced at OD_600_ 0.7 with aTc, IPTG, or both accordingly. After the induction the cultures were grown for 4 hours, 2 mL samples were collected, centrifuged at 14,000 rpm, the supernatant was discarded, and the cell pellets were stored at −80°C. The cell pellets were resuspended with 200 µL acetone and incubated at 55°C for 15 min in the dark followed by centrifugation at 14,000 rpm, then the supernatant was filtered through a 0.22 μm pore-size nylon membrane for UHPLC analysis.

### Bioluminescence assay

*E. coli* or *P. putida* overnight cultures supplemented with kanamycin (LB, 37°C with shaking, 220 rpm) were diluted 1/200 in fresh LB in 96-well black microplates (Corning Incorporated, NY) shaking at 355 cycles per minute (cpm) and induced after three hours of growth with 0.2/ 5 mM IPTG and 0.1/1.0 µg/mL aTc accordingly. Bioluminescence (gain: 100) and cell density (OD_600_) were monitored every 20 minutes with the BioTek Synergy H1 Hybrid Multi-Mode Reader instrument for 20 hours.

## RESULTS AND DISCUSSION

### Development of the dual expression system

Duet vectors are available for *E. coli* and *Pseudomonas fluorescens*, but they are solely inducible by either IPTG or arabinose [26, 39], whereas dual-inducible duet expression vectors have been developed specifically for *E. coli* [28], *Corynebacterium glutamicum* [40] and *P. putida.* Our previous results indicated that the *σ*^70^V2TcR and σ^70^V3LacI expression systems exhibited tight regulation and robust induction in *E. coli* in the presence of anhydrotetracycline (aTc) and IPTG, respectively, while in *P. putida* and *V. natriegens*, the *σ*^70^V2TcR and *σ*^70^V2LacI promoters exhibited enhanced performance as inducible promoters when compared to the original *lac* and *tet* promoters [23]. These newly engineered promoters not only yielded higher amounts of recombinant proteins upon induction but also maintained lower expression levels in the absence of the inducers, thereby providing a more controlled and efficient production system [23]. We developed the dual expression system using the pJH0204 vector as a backbone, which contains the pColE1 origin or replication that replicates in *E. coli* and *V. natriegens*, and the *attB* sites specific for the bxb1 recombinase, to integrate it into the chromosome of *P. putida* [41]. Each transcriptional unit of the dual expression system was insulated by a terminator [42]. The transcriptional regulators *tetR* and *lacI* with their corresponding constitutive promoters pJ23119 and placI were positioned in-between the inducible promoters *σ^70^*V2Tc and *σ^70^*V2Lac or *σ^70^*V3Lac (*σ^70^*V2/3Lac) and in opposite direction to block any undesired transcription triggered by the constitutive promoters. Additionally, two different multiple cloning sites were placed after the inducible promoters *σ^70^*V2Tc and *σ^70^*V2/3Lac to facilitate the incorporation of target genes (Fig. 1).

**Figure 1.**
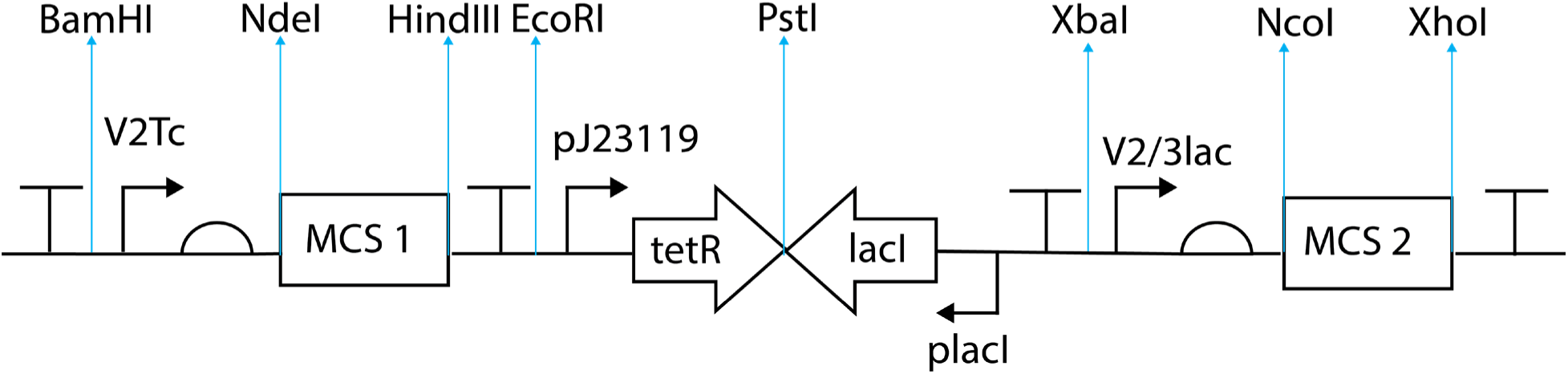
Design of the dual expression systems σ^70^V2TcR-V2LacI and σ^70^V2TcR-V3LacI

The dual expression system was assessed in *E. coli*, *P. putida*, and *V. natriegens* to evaluate: i) independent control of two different fluorescent proteins, ii) potential synergy (expression in excess of additive transcription) of both promoters when expressing the same protein, and iii) control of the biosynthetic gene cluster (BGC) responsible for the biosynthesis of lycopene and β-carotene. Expression of various recombinant proteins was achieved by placing *sfGFP* under the control of *σ^70^*V2TcR and *mCardinal* under the control of *σ^70^*V2LacI and V3LacI (Fig. 2C). Synergy was gauged using the reporter system *mCardinal*, which was incorporated into both inducible promoters of the dual expression system (Fig. 3B). Finally, production of the terpenoids lycopene and β-carotene by the dual expression system was achieved by the assembly of the *crtEBIY* genes from *Pantoea ananatis* [11], which were synthesized with the codon usage for *E. coli.* The *crtEBI* (LYC operon) was assembled in the pUC19 vector via Gibson assembly, adapting the ribosome binding site (RBS) of J23119 [43] to *crtB* and the RBS pJL1 [44] to *crtI,* then the synthetic construct was transferred to the *σ^70^*V2TcR expression system to evaluate lycopene production (Fig. 4A). β-carotene production was evaluated with the incorporation of the *crtY* gene with the J23119 RBS to *σ^70^*V2TcR-*crtEBI* yielding *σ^70^*V2TcR-*crtEBIY* (Fig. 4B). The dual control of the *crtEBI* operon together with the *crtY* gene to produce β-carotene was achieved by adapting the *crtEBI* to the σ^70^V2TcR expression system and the *crtY* gene to σ^70^V2/3LacI expression system present in the dual vector (Fig. 4E).

**Figure 2.**
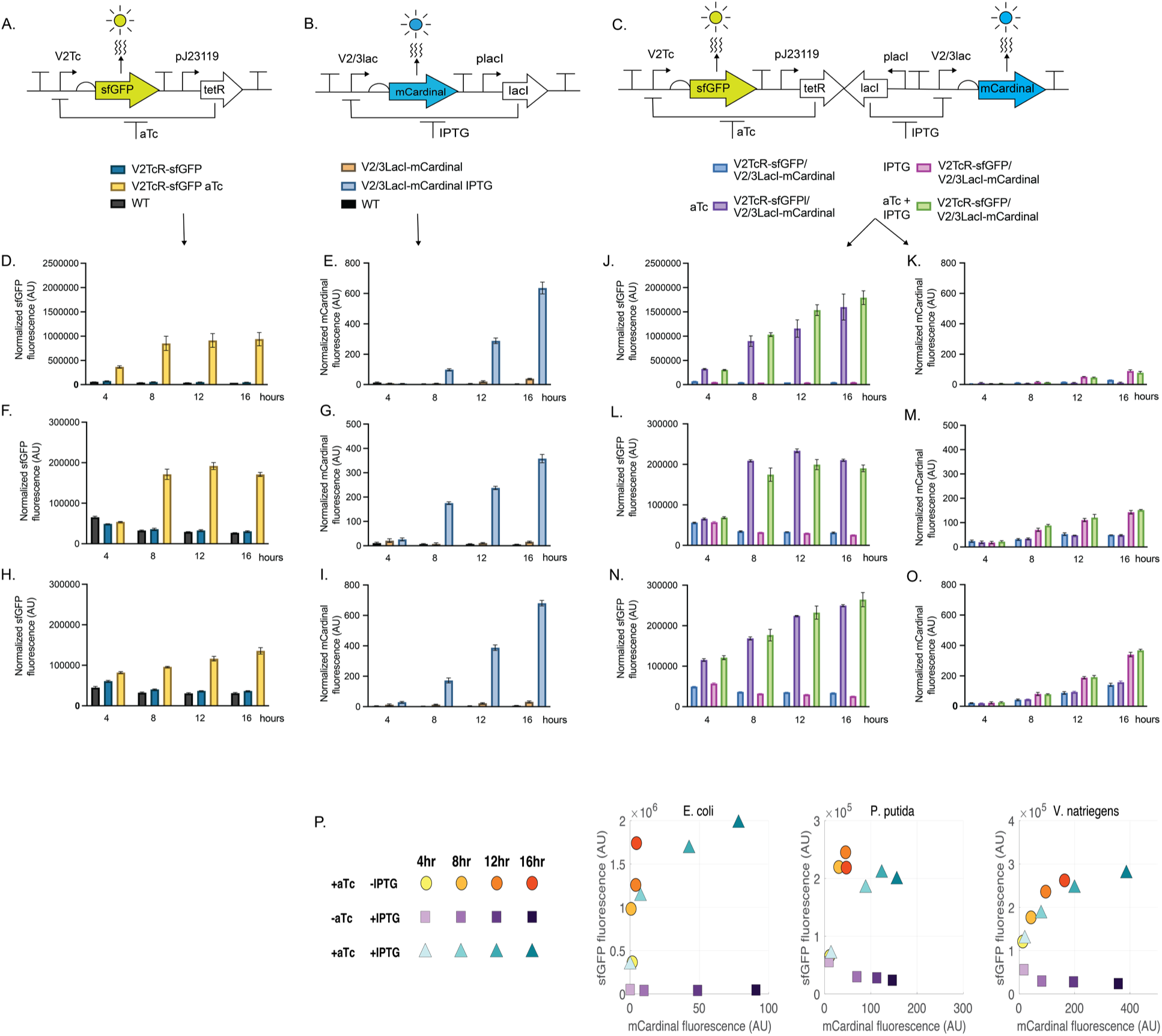
*sfGFP* and *mCardinal* production in *E. coli*, *P. putida*, and *V. natriegens* containing the single and dual expression systems. For all samples, the fluorescence mean of *sfGFP* signal (excitation 485, emission 510) and *mCardinal* signal (excitation 605, emission 659) was normalized by the cell density (OD_600_). Gene circuits of **A.** σ^70^V2TcR-*sfGFP* **B.** σ^70^V2/3LacI-*mCardinal* & **C.** σ^70^V2TcR-*sfGFP*/σ^70^V3LacI-*mCardinal*. Normalized *sfGFP* emission produced by σ^70^V2TcR-*sfGFP* in **D.** *E. coli* **F.** *P. putida* & **H.** *V. natriegens*. Normalized *mCardinal* emission produced by σ^70^V2/3LacI-*mCardinal* in **E.** *E. coli* **G.** *P. putida* & **I.** *V. natriegens*. Normalized *sfGFP* and *mCardinal* emission produced by the dual expression system σ^70^V2TcR-*sfGFP*/σ^70^V2/3LacI-*mCardinal* in **J.K** *E. coli* **L.M.** *P. putida* & **N.O.** *V. natriegens*. (Error bars are +/- S.D., n=3. **P.** Matrix display of both *sfGFP* and *mCardinal* production of individual and concurrent inductions of the dual expression systems.

**Figure 3.**
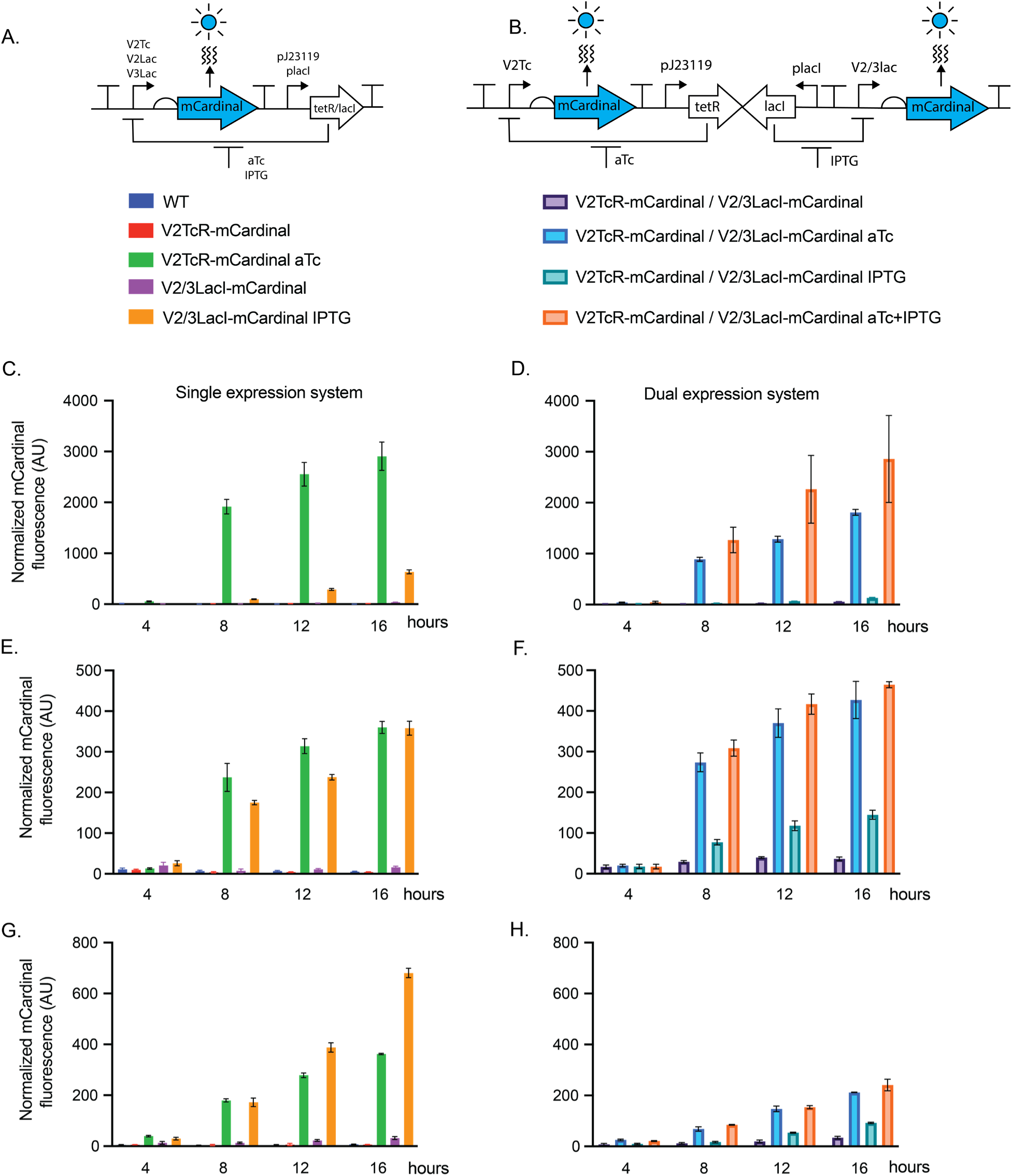
Assessing potential synergy between *lac* and *tet* promoters. Gene circuits of the single expression system **A.** σ^70^V2TcR-*mCardinal*, σ^70^V2LacI-*mCardinal* or σ^70^V3LacI-*mCardinal* & the dual expression system **B.** *σ^70^*V2TcR-*mCardinal*-V2/3LacI-*mCardinal*. Time course of *mCardinal* production from the single expression systems in **C.** *E. coli* **E.** *P. putida* & **G.** *V. natriegens* Vs *mCardinal* production in the double expression system in **D.** *E. coli* **F**. *P. putida* & **H.** *V. natriegens*. For all samples, the fluorescence mean of *mCardinal* signal (excitation 605 nm, emission 659 nm) was normalized by the cell density (OD_600_). Error bars are +/- S.D., n=3.

**Figure 4.**
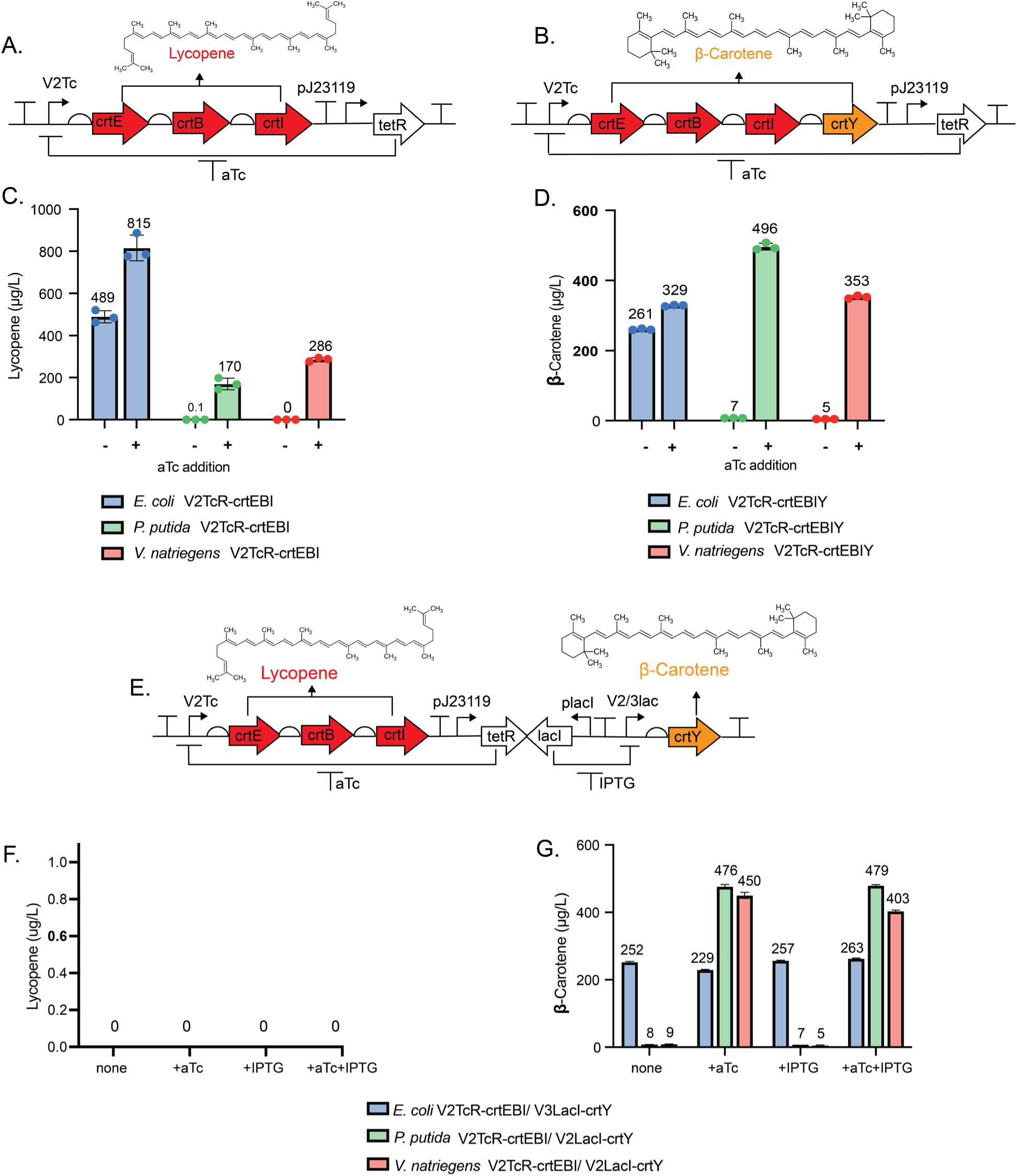
Production of lycopene and β-carotene in *E. coli, P. putida,* and *V. natriegens* with the single and dual expression systems. Gene circuit of **A.** σ^70^V2TcR-*crtEBI* and **B.** σ^70^V2TcR-*crtEBIY*. Production of **C.** lycopene and **D.** β-carotene by the single σ^70^V2TcR expression system in *E. coli, P. putida* and *V. natriegens*. Gene circuit of **E.** the dual expression system σ^70^V2TcR-*crtEBI*-V2/3LacI-*crtY*. Production of **F.** lycopene and **G.** β-carotene by the dual expression system in *E. coli, P. putida* and *V. natriegens*. Cultures were induced with aTc, IPTG or both inducers at OD_600_. 0.7 for 4 hours and terpenoids titers were measured by UHPLC. Error bars are +/- S.D., n=3.

### Independent and concurrent control of two fluorescent proteins with the dual expression system

We have previously demonstrated the strength and portability of the *σ^70^*V2TcR, *σ^70^*V2LacI and *σ^70^*V3LacI promoters separately in *E. coli, P. putida*, and *V. natriegens* [23]. To test the independent and concurrent regulation of two proteins simultaneously from these promoters, we introduced two different reporter proteins into the dual-inducible *σ^70^*V2TcR/V2LacI and *σ^70^*V2TcR/V3LacI expression systems (Fig. 2C) and assessed their production in *E. coli*, *P. putida*, and *V. natriegens*. Specifically, the transcriptional unit V2TcR controlled the expression of *sfGFP*, while the V2LacI and V3LacI unit regulated *mCardinal* production (Fig. 2C). This configuration was chosen because the *σ^70^*V2TcR expression system exhibited similar performance across the three bacterial species under study [23]. However, the *σ^70^*V2LacI system showed strong recombinant protein production in *E. coli* even in the uninduced state, undermining the benefits of an inducible system [23]. As a result, the *σ^70^*V3LacI system was selected for *E. coli*, while the *σ^70^*V2LacI system was further evaluated in *P. putida* and *V. natriegens*. Due to the endogenous autofluorescence in the yellow-green spectrum of these bacteria [23], which could interfere with *sfGFP* measurement, the *σ^70^*V2TcR promoter was selected to regulate *sfGFP* expression. This promoter provides tighter control in the OFF state, allowing for a more accurate and sensitive estimation of reporter system production. On the other hand, the *σ^70^*VLacI systems, which tend to leak [23], control the expression of the far-red fluorescent protein *mCardinal*. Importantly, at this wavelength (excitation 604 nm, emission 659 nm), the cells do not produce endogenous fluorescence [45].

To create a direct comparison, the induction of the individual expression systems *σ^70^*V2TcR-*sfGFP* was measured in *E. coli* (Fig. 2D), *P. putida* (Fig. 2F) and *V. natriegens* (Fig. 2H), while the *σ^70^*V3LacI-*mCardinal* was evaluated in *E. coli* (Fig. 2E) and the *σ^70^*V2LacI-*mCardinal* in *P. putida* (Fig. 2G) and *V. natriegens* (Fig. 2I). The dual-inducible *σ^70^*V2TcR/V2LacI and *σ^70^*V2TcR/V3LacI expression systems were also measured in the same species (Fig. 2J-O). Under the control of *σ^70^*V2TcR, *sfGFP* production was higher in the dual system by 1.2-fold in *E. coli* and *P. putida*, and 1.7-fold in *V. natriegens* at 16 hours of induction (Fig. 2J, L, N). However, an opposite trend was observed in the dual expression system for production of *mCardinal* under the control of *σ^70^*V2/3LacI, which was significantly reduced, resulting in 6.2-fold, 2.4-fold, and 2.0-fold less red emission in *E. coli*, *P. putida*, and *V. natriegens*, respectively (Fig. 2K, M, O).

Interestingly, when the cultures containing the dual expression system were co-induced with aTc and IPTG, both green and red fluorescence reached comparable levels compared to cultures induced with only one inducer. So, while the maximum expression from each promoter was distinct from the maxima achieved in the single promoter systems, concurrent induction of both promoters did not further impact their transcriptional output. This suggests that the promoters (*σ^70^*V2TcR and *σ^70^*V2/3LacI) are minimally affected by the activation of the other, reaching almost their maximum activity either upon sole activation or concurrent activation (Fig. 2J-O). The three bacterial species investigated, each containing the dual expression system, did not display detectable green fluorescence in the uninduced state compared to their wild type counterparts, indicating tight regulation of the *σ^70^*V2TcR expression system (Fig. 2D, F & H). However, as we previously observed [23], some *mCardinal* leakage was observed originating from the *σ^70^*V2LacI-*mCardinal* (Fig. 2E, G & I).

The dual expression system constitutively expresses the transcriptional regulators *TetR* and *LacI*, both of which have been reported to exhibit toxicity at high concentrations [46–48]. Each sensor requires cellular resources (for example, ribosomes) to function, and the activation of one sensor can influence another indirectly due to resource competition [49]. We anticipated a lower performance for both promoters in the dual expression system, due to the metabolic burden imposed by the simultaneous presence of *TetR* and *LacI*. However, this scenario was only observed with *σ^70^*V2/3LacI, which exhibited reduced production of *mCardinal*, which was particularly pronounced in *E. coli* (Fig. 2K, M & O), thus suggesting that the presence of both transcriptional regulators might affect other cellular processes that indirectly affect the strength of the *σ^70^*Lac promoter. Conversely, the levels of *sfGFP* appeared to be enhanced by *σ^70^*V2TcR in the dual expression system. One hypothesis for this observation is that our transcriptional systems are stressing the cells with their metabolic load. To mitigate the adverse effect of stresses, some cells produce endogenous fluorophores and/or metabolic byproducts increasing the emission of green light [50]. For instance, *P. putida* secretes pyoverdine, a soluble fluorescent yellow-green pigment, under iron-limiting conditions [51]. We predict that *σ^70^*V2TcR in the dual expression system did not in fact display increased *sfGFP* production. Instead, it is possible that the presence of endogenous molecules interfering in the green spectrum [52, 53], not originating from the recombinant protein, amplified the *sfGFP* signal due to activation of cellular stress responses. Similarly, we would predict that the cells, in addition to increasing their autofluorescence, would display changes in growth that reflect this additional metabolic load.

To corroborate these hypotheses, we examined the growth rate of the parent *E. coli* strain (DH10B) and the recombinant strains carrying the dual expression system. Addition of IPTG or aTc shows minimal effect in the cell growth (Fig. S1A-B), however the combined exposure to both inducers imparts a modest effect on the growth rate (Fig. S1C). A major impact on growth is observed when the dual expression system is incorporated into *E. coli* in pColE1 plasmid, reaching the beginning of the exponential phase 4 hours after the wild type (Fig. S1D). Addition of IPTG does not delay the growth rate further in the recombinant strains (Fig. S1E), however, aTc (Fig. S1F) or aTc with IPTG (Fig. S1G) supplementation prolongs the exponential phase.

The emission of green fluorescence was assessed to identify any potential endogenous molecules or proteins that might interfere with the readings of the recombinant sfGFP. The wild-type DH10B strain does not produce any additional green fluorescence when exposed to IPTG, aTc, or a combination of both (Fig. S2A-C). However, the recombinant strain with the dual expression system, without *sfGFP* and containing only mCardinal, exhibits significantly higher green fluorescence compared to the wild type, showing a maximum 5-fold increase after 4 hours of growth (Fig. S2D). Induction with IPTG does not increase green fluorescence emission (Fig. S2E). However, the addition of aTc (Fig. S2F) and aTc combined with IPTG (Fig. S2G) results in approximately a 2-fold increase in green emission compared to the uninduced state. It is tempting to conclude that perhaps aTc is itself the source of the fluorescence, however these data are inconsistent with that conclusion because additional green fluorescence is not observed in the parent DH10B strain when aTc is added. Therefore, the results are consistent with the idea that the cells carrying the dual expression system are generating a fluorescent metabolite of some kind, which correlates with a growth defect.

### Evaluating synergistic potential and metabolic cost of the dual expression system

The combinatorial use of two promoters that control transcription of the same gene, or double promoter expression systems, have the potential to increase the titer of recombinant protein production [29, 54]. However, from our first dual expression system experiments, we anticipated a metabolic cost to the cell (Fig. S3). We observed that while *sfGFP* fluorescence from the *σ^70^*V2Tc promoter increased relative to a single expression system, *mCardinal* fluorescence from the *σ^70^*V2/3Lac promoters was significantly decreased. If, as we hypothesized, the increased *sfGFP* was an artifact of cellular stress caused by the combination of metabolic load and aTc exposure, then one would predict that driving the same protein from both promoters would have no synergistic effect on expression over a single promoter system.

Previous results indicated that recombinant protein production by *σ^70^*V2TcR expressed in pJH0204 (pColE1 derived plasmid with 30 copies per cell) was enhanced by increasing the copy number by adapting the expression system to pUC19 vector (250 copies per cell). However, the toxicity associated with the transcriptional regulator *LacI* prevented the evaluation of the *σ^70^*V2/3LacI expression system in a high copy plasmid like pUC19 [23]. Hence, to test the synergistic potential of the duet expression system *σ^70^*V2TcR/V3LacI, a version was made where both promoters regulate *mCardinal* production (Fig. 3B). The *E. coli* strain harboring the duet expression system *σ^70^*V2TcR/V3LacI exhibited 2-fold and 4-fold lower red fluorescence compared to strains containing *σ^70^*V2TcR and *σ^70^*V3LacI individually when induced with aTc and IPTG, respectively (Fig. 3C, D). This observation supports the hypothesis that the increased green fluorescence signal in the *σ^70^*V2TcR/V3LacI was an artifact of some kind, possibly a product of cellular stress. Synergy was observed by the addition of IPTG and aTc to the *σ^70^*V2TcR/V3LacI *E. coli* strain improving *mCardinal* production by 1.5-fold against the induction obtained with aTc alone (Fig. 3D). However, the *mCardinal* production in the duet system reached about the same total amount of *mCardinal* as the *σ^70^*V2TcR expression system alone (Fig. 3C, D). These results indicate that the tandem version of the synthetic promoters *σ^70^*V2TcR/V3LacI does not offer a real synergy in *E. coli*, but rather an alternative to co-express different genes at high levels using different inducers.

In *P. putida* the *σ^70^*V2TcR expression system exhibited similar *mCardinal* performance in the single and dual version (Fig. 3E & F). Conversely, induction of the dual system with IPTG resulted in a 3-fold decrease relative to the *σ^70^*V2lacI system alone (Fig. 3E, F). The simultaneous addition of both inducers (aTc and IPTG) increased *mCardinal* expression by 1.1-fold over the *σ^70^*V2TcR and *σ^70^*V2LacI expression systems in the dual system, demonstrating a minimal, statistically insignificant synergistic effect in this host. The duet expression system *σ^70^*V2TcR/V2LacI markedly reduced the efficiency of the synthetic promoters in *V. natriegens* compared to their single expression system counterparts, with reductions of 12-fold and 2-fold for *σ^70^*V2LacI and *σ^70^*V2TcR after induction with IPTG and aTc, respectively (Fig. 3G & H). Co-induction of the dual expression system with IPTG and aTc did not yield a synergistic effect in *mCardinal* production, but achieved similar yields as obtained by induction with aTc alone (Fig. 3H).

This information collectively demonstrates that synergy cannot be attained with the dual expression system in the three microorganisms under study. In *E. coli* and *V. natriegens*, the pJH0204 plasmid containing the pColE1 origin of replication was used, rendering 30 copies per cell in *E. coli* and 300 copies per cell in *V. natriegens*. [14, 55]. On the contrary, the duet expression system appears to attenuate the activity of each promoter. Rewiring the cell metabolism imposes additional stress which affects protein synthesis and growth rate [56]. Consequently, the constitutive expression of the transcriptional regulators *TetR* and *LacI*, as well as competition for the *σ^70^* factor by both promoters, may pose a significant metabolic burden. This triggers energetic inefficiency within the cell, leading to undesirable physiological changes and placing hidden constraints on host productivity [57]. In *P. putida*, the dual expression system was directly integrated into the chromosome, thereby reducing the quantities of the transcriptional regulators *TetR* and *LacI* and thus lowering the metabolic cost of the synthetic expression systems. This resulted in similar *mCardinal* production with *σ^70^*V2TcR, while the *σ^70^*V2LacI expression system showed reduced expression and a lack of synergy.

### Production of lycopene and β-carotene in *E. coli*, *P. putida* and *V. natriegens* using the σ^70^V2TcR single expression system

To assess the capability of the dual expression system to produce natural products, we tested the ability of the *σ^70^*V2TcR expression system to control the synthesis of multiple genes in a BGC. To do so, we evaluated the production of lycopene and β-carotene. Lycopene production has already been demonstrated in *E. coli* and *P. putida* through heterologous expression of the lycopene-producing operon (LYC), which contains the *crtEBI* genes from *Pantoea ananatis* [11]. Lycopene serves as the biosynthetic precursor of β-carotene. Therefore, the expression of the LYC operon together with *crtY* leads to β-carotene production. This process has also been demonstrated in *E. coli* and *V. natriegens* [13, 58]. The LYC and *crtEBIY* operons were incorporated into the *σ^70^*V2TcR expression system (Fig. 4A, B) and the production of lycopene and β-carotene was quantified using UHPLC [37] after 4 hours of induction with aTc. In *E. coli*, production of both lycopene and β-carotene was achieved. However, the induced culture only yielded 1.7-fold more lycopene and 1.2-fold more β-carotene compared to the uninduced culture when induced for four hours (Fig. 4C, D). A decrease in lycopene production was noted with longer induction times which may be due to the documented toxicity of high lycopene production in *E. coli* that inhibits cell growth and productivity, thereby decreasing titer per volume [59]. The results demonstrate that the tight control of the *σ^70^*V2TcR system is still unable to completely repress the expression of the *crtEBI* and *crtEBIY* BGCs in *E. coli*. A similar observation has been previously published, where lycopene biosynthesis in *E. coli* was controlled using the pET expression system, and also showed significant leakage, with production reaching only 1.1-fold more after 40 hours of induction than the uninduced culture [38]. Clearly, the tight regulation of terpenoid biosynthesis in *E. coli* remains a significant challenge.

The *σ^70^*V2TcR expression system produced a maximum 0.8 mg/L of lycopene in *E. coli* cultivated in a culture tube after only 7 hours of growth (4 hours with the inducer aTc) (Fig. 4C), and a maximum of 0.46 mg/L β-carotene (Fig. 4D). Notably, no media optimizations were performed, nor any strain engineering beyond the addition of the *crtEBI* or *crtEBIY* BGCs. While higher titers of lycopene and β-carotene have been reported in *E. coli* by rewiring its metabolic network, and utilizing optimal media and bioreactors [38], our goal here was to optimize control, rather than titer. Still, the titers obtained with the *σ^70^*V2TcR expression are comparable to those generated in some previous reports, even in the absence of further optimizations [11, 13, 38, 58]. These titers could be further increased through media and strain optimization in future studies.

In recombinant strains of *P. putida* and *V. natriegens* containing the *σ^70^*V2TcR-*crtEBI* and *σ^70^*V2TcR-*crtEBIY* transcriptional units, little to no lycopene or β-carotene production was detected without induction. In both organisms, induction of the BGC by aTc led to the robust production of the terpenoids (Fig. 4C, D). *P. putida* was reported to produce 1.233 µg/L of lycopene after 24 hours of growth with the expression of *crtEBI* from pSEVA421 vector, in the absence of further strain engineering [11]. With the *σ^70^*V2TcR expression system, lycopene production reached 170 µg/L after only 4 hours of induction (Fig. 4C), marking a substantial increase in reported titer. Additionally, β-carotene production by *P. putida* reached 496 µg/L, the highest titer among the three species in our study (Fig. 4D). *V. natriegens* was reported to produce 500 µg/L of β-carotene cultivated in the rich medium LBv2 for 20 hours [13], while in the *σ^70^*V2TcR expression system achieved 286 µg/L of lycopene (Fig. 4C) and 353 µg/L of β-carotene (Fig. 4D) in the standard LB medium and four hours of induction with no particular optimization. These titers achieved here suggest that significantly increased titers would be possible in the future with the combination of our expression system and other optimizations. The overall results demonstrate that the *σ^70^*V2TcR expression system can be efficiently used to drive production of BGCs in *E. coli*, *P. putida* and *V. natriegens*.

### Coordinated expression of the lycopene and β-carotene BGCs with the dual expression system

Natural products are often encoded by BGCs, and the ability to activate each gene at different time points with different inducers can be advantageous, particularly when the genes are toxic or the metabolites produced by the BGC affect the growth of the heterologous host [60]. Such control may also be useful in industrial bioprocess settings, permitting independent optimization and control of different enzymatic steps in a pathway. In this study, we have demonstrated that the *σ^70^*V2TcR expression system can effectively activate the production of lycopene and β-carotene. As a next step, we attempted to exert control of β-carotene at different points in the production pathway using two different inducers, aTc and IPTG. The LYC operon was integrated into the *σ^70^*V2TcR expression system, while the *crtY* gene was controlled by the *σ^70^*V2/3LacI expression system (Fig. 4E). The plasmid p*σ^70^*V2TcR-*crtEBI*/V3LacI-*crtY* was transformed into *E. coli*, and the plasmid p*σ^70^*V2TcR-*crtEBI*/V2lacI-*crtY* was transformed into *P. putida* and *V. natriegens*. In all three bacterial species, no lycopene accumulation was observed upon sole induction by aTc despite the *crtEBI* genes being controlled by the *σ^70^*V2TcR expression system (Fig. 4F). This result in *E. coli* can be explained by the leakage of the *crtY* under the control of the synthetic promoter *σ^70^*V3LacI. The high processivity of the lycopene cyclase CrtY to metabolize lycopene directly into β-carotene requires tighter regulation of the promoter to completely repress this gene, and the *σ^70^*V3LacI expression systems lacks this control. On the other hand, while the *σ^70^*V2TcR-*crtEBI* system showed no measurable leakage in *P. putida* and *V. natriegens* using fluorescent reporters, we previously showed that the *σ^70^*V2lacI expression system does leak in these strains [23]. Consequently, the lycopene produced after the addition of aTc was directly processed into β-carotene by the CrtY present in the cells due to leakage of *σ^70^*V2LacI.

Production of β-carotene in *E. coli* was observed regardless of the absence or presence of the inducers aTc and IPTG (Fig. 4G), demonstrating that the dual *σ^70^*V2TcR/ *σ^70^*V3LacI expression system would require tighter regulation in the OFF state to effectively downregulate the target gene in the absence of the inducer. We believe the leaky transcription we are observing in the lycopene and β-carotene BGCs, which appears more severe than was observed for similar promoter designs expressing fluorescent proteins, is not due to an increase in leaky transcription from these promoters under carotenoid biosynthesis conditions, but rather to the processivity of enzymes versus fluorescent proteins. In *P. putida* and *V. natriegens*, production of β-carotene was only achieved when the cultures were induced with aTc or with aTc + IPTG (Fig. 4G). IPTG induction alone did not lead to β-carotene production because the *crtEBI* genes were not transcribed by the *σ^70^*V2TcR expression system. Addition of aTc resulted in β-carotene levels similar to the ones obtained when the cells were induced with aTc + IPTG (Fig. 4G), thus indicating that the minimal leakage of the *σ^70^*V2lacI-*crtY* in *P. putida* and *V. natriegens* is sufficient to metabolize all lycopene produced by the *σ^70^*V2TcR-*crtEBI* immediately into β-carotene upon induction by aTc. The varying yields of β-carotene between *E. coli*, *P. putida*, and *V. natriegens* highlight the differences in their suitability as hosts for controlled natural product production.

### Adaptation of toehold switch to the synthetic *σ^70^*expression systems and fold-change evaluation with the reporter system *mCardinal*

The assessment of *σ^70^* expression systems using the reporter *mCardinal* suggested that the *σ^70^*V2TcR promoter displays tight regulation, leading to undetectable levels of red fluorescence in the uninduced state [23]. However, when the lycopene and β-carotene operons were incorporated into this synthetic promoter, the terpenoids were produced by *E. coli* in the uninduced state, indicating that the promoter was not completely repressed. The processivity of enzymes, unlike static fluorescent proteins, amplifies the effect of leaky transcription. In the *σ^70^* V2/3LacI expression systems the leakage was even more evident, as we were unable to repress the production of CrtY in the dual expression system sufficiently to permit the intermediate accumulation of lycopene (Fig. 4F). Maintaining low basal expression levels in the absence of the inducer is crucial to minimize unwanted effects on cell growth and to prevent potential damage caused by the target protein, including issues like self-aggregation, inclusion body formation [61], or as evidenced in this study, uncontrolled biosynthesis of a downstream product. If independent control of steps within a multistep biosynthetic pathway is to be achieved, it will require significant improvements in the control of inducible promoter systems.

One solution to this problem involves the adaptation of RNA-based gene regulation to reduce the impact of basal expression [32, 33]. The current design of the synthetic *σ^70^* expression system comprises transcriptional control performed by the transcriptional regulators TetR and LacI. Introducing an additional translational control mechanism, such as a toehold switch (THS), alongside the existing transcriptional control, could enhance the precision of the *σ^70^* expression systems. This added layer of regulation could provide finer control over gene expression, potentially reducing leakiness and improving inducibility. To this end, the *σ^70^*V2TcR and *σ^70^*V2lacI expression systems were modified with the incorporation of the three components required to add toehold-mediated translational control: the THS trigger, the THS target and the cognate ribosome binding site (RBS) [34]. For the *σ^70^*V2TcR expression system, two *σ^70^*V2Tc promoters were incorporated into the system, the first *σ^70^*V2Tc promoter regulates the transcription of the THS trigger, while the second promoter is fused to the THS target, which masks the RBS (Fig. 5A). The *σ^70^*V2lacI follows the same architecture, except the THS trigger is regulated by the V3Lac promoter due its reduced leakage compared to the V2Lac, which controls the THS target that hides the RBS (Fig. 5B). This arrangement combines transcriptional and translation regulation, therefore, the absence of the inducers (aTc or IPTG) results in both transcriptional repression (via the transcriptional regulators *TetR* and *LacI*) and translational repression (via the THS trigger) of the target gene (*mCardinal*). Upon addition of the inducers, the constitutively expressed transcriptional regulators are removed from the synthetic *σ^70^* promoters, activating the transcription of the THS trigger, which binds to the THS target inducing the conformational change that relieves the translational repression of *mCardinal* resulting in its subsequent expression.

**Figure 5.**
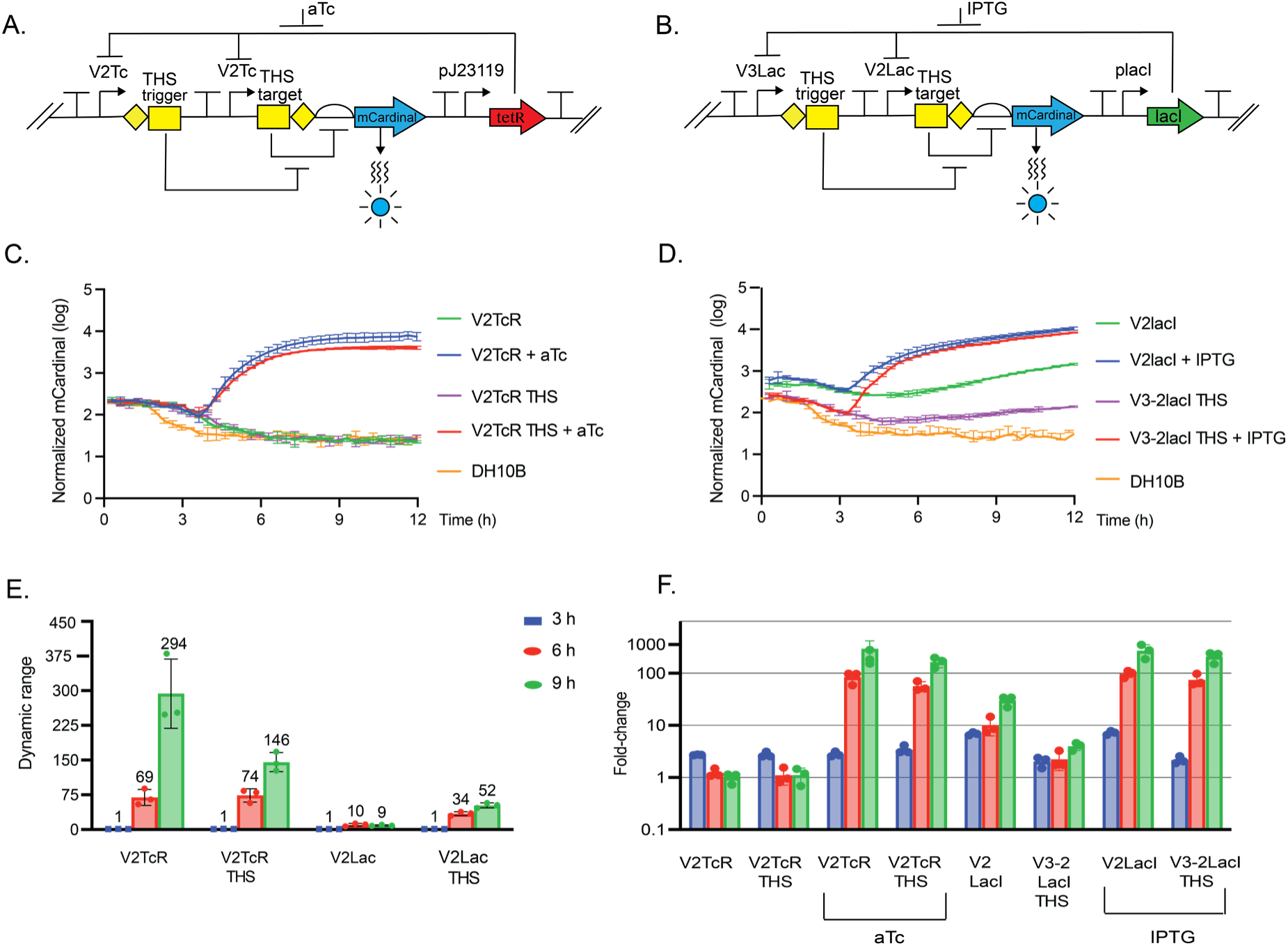
Comparison of the σ^70^ expression systems to the THS versions controlling the expression of the reporter system *mCardinal* in *E. coli*. Architecture of the synthetic gene circuits with the toehold switches (THSs) in **A.** σ^70^V2TcR & **B.** the σ^70^V3-2LacI system. **C**. Time course of *mCardinal* production by pσ^70^V2TcR THS. **D**. Time course of *mCardinal* production by σ^70^V3-2LacI THS. **E.** Difference in *mCardinal* production between the ON and OFF states. **F**. Fold-change difference in the red-light emission of the recombinant cells vs the wild type DH10B. The cultures were induced after 3 hours of growth with aTc or IPTG accordingly. For all samples, the fluorescence mean of *mCardinal* signal (excitation 605 nm, emission 659 nm) was normalized by the cell density (OD_600_). Error bars are +/- S.D., n=3.

The comparison between the uninduced and induced states of the *σ^70^*V2TcR expression system with and without the toehold switch revealed very similar induction dynamics (Fig. 5C). It was previously determined that the leakiness of *σ^70^*V2TcR could not be observed with a fluorescent reporter, thus no change in the performance of this reporter was expected. The system may exhibit leakage even with the addition of the THS, but it would not be detectable in this assay. The addition of aTc induced *mCardinal* production by promoters with and without the THS. However, the promoter with the THS exhibited a 2-fold reduction in the dynamic range compared to the promoter without the switch, although it still maintained a satisfactory level of recombinant protein production (Fig. 5C, E, F). On the other hand, the addition of the THS had a significant effect on suppressing leaky transcription from the *σ^70^*V2lacI expression system (Fig. 5D). The dynamic range in the *σ^70^*V3-2lacI THS system improved by 5.7-fold, but still, leaky transcription was significant. However, the THS did reduce the leakiness by 7-fold compared to the original version (Fig. 5E). Importantly, *mCardinal* production after IPTG induction reached similar levels in both *σ^70^*V2lacI expressions, with the variant lacking the THS producing only 1.2-fold more *mCardinal* fluorescence (Fig. 5D).

These results demonstrate that THSs enhance the regulation of gene expression by modulating translation initiation (Fig. 5E, F). This feature is particularly effective in inducible expression systems prone to leakage. However, while fluorescent proteins like *mCardinal* offer valuable insights, they might not capture the subtle nature of basal transcription. For instance, the evaluation of the *σ^70^*V2TcR expression system using fluorescent proteins indicated complete lack of fluorescent protein synthesis in the OFF state when uninduced. However, the results presented for the production of lycopene in the dual expression system revealed that the promoter leaks to some extent. Therefore, to truly assess any basal transcriptional activity, the use of enzymatic reporter systems will be necessary, with higher sensitivity for accurately evaluating gene expression dynamics.

### Evaluation of the *σ^70^* expression systems using luciferase as a reporter

The selection of the red fluorescent protein *mCardinal* to estimate the efficiency of the *σ^70^* expression systems was a strategic choice, to circumvent intrinsic issues associated with *sfGFP* and the potential interference of green autofluorescence from cells, which could lead to inaccurate quantification of the recombinant protein produced by the synthetic promoters [23]. However, upon assessing the synthetic promoters for terpenoid production, it became evident that the *σ^70^*V2TcR expression system does display leaky transcription, a phenomenon not discernible with *mCardinal* as the reporter. The integration of the THS into *σ^70^*V2lacI effectively mitigated basal expression. However, this effect was not evident in THS-*σ^70^*V2TcR-*mCardinal*, as undetectable fluorescence (undetectable above the background autofluorescence) in the OFF state in this expression system was achieved even without the THS. The terpenoids lycopene and β-carotene, aside from their nutraceutical properties, can serve as reporter systems due to their ability to induce a color change in the cell biomass upon production [62]. Indeed, the biosynthesis of these pigments was instrumental in detecting the leakage of the *σ^70^*V2TcR system. However, accurate estimation of terpenoid yields necessitates laborious compound isolation techniques and complex measurements. Consequently, their use as a reporter system may not be the ideal choice for evaluating the metrics of an inducible system. To comprehensively grasp the behavior of the synthetic *σ^70^*expression systems and address the challenges posed by the limited sensitivity of *mCardinal* and the intricate evaluation of terpenoid production, we opted to employ luciferase as a reporter.

The bacterial luciferase, encoded by the lux operon (*luxCDABE*), initiates the emission of bioluminescence upon its expression, without requiring the addition of an external substrate [63]. To amplify the detection of leaky basal transcription, the luciferase reporter was integrated into the *σ^70^* expression systems and their corresponding THS variants in the pColE1-containing plasmid pJH2024 (Fig. 6A - D). We expected these constructs would allow us to better measure leaky transcription, particularly in the *σ^70^*V2TcR system where the uninduced state appears completely OFF when using *mCardinal*, yet shows activity with lycopene and β-carotene (Fig. 4C, D). The time course evaluation of luciferase production permitted the visualization of the leakage of the *σ^70^*V2TcR, and also confirmed that the THS version of this promoter diminished the leakage and amplified the signal of bioluminescence in *E. coli* (Fig. 6E, G, I). Interestingly, bioluminescence production with the THS tends to diminish slightly over time, suggesting that translational regulation favors the repression of the lux operon after the mid-log phase of growth (Fig. S4). In contrast, the original *σ^70^*V2TcR system maintains its pronounced leakage over time (Fig. 6E). The *σ^70^*V2lacI variant, exhibited high basal leakage under both induced and uninduced conditions, with only a modest improvement in induction by the integration of the THS (Fig. 6F). Interestingly, induction with IPTG in the *σ^70^*V2lacI variant slightly suppressed bioluminescence production (Fig. 6F, blue bars versus red bars). This phenomenon may be attributed to the potency of this promoter, whose activation could exceed the metabolic capacity of the cells.

**Figure 6.**
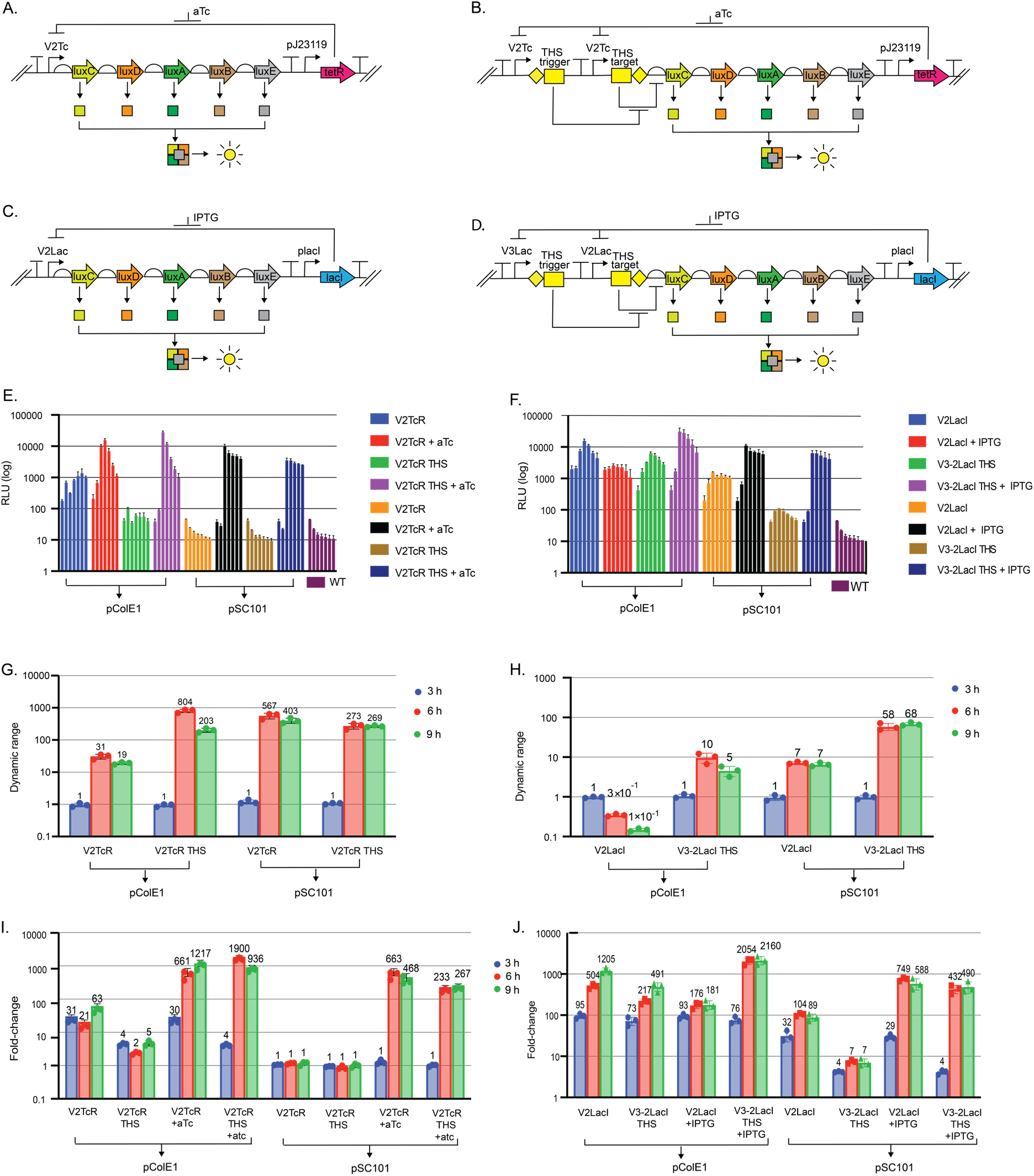
Comparison of the σ^70^ expression systems to the toehold versions controlling the expression of the lux operon in *E. coli* using pCOLE1 and pSC101 derived plasmids. Architecture of the synthetic gene circuits controlling the *luxCDABE* genes with and without the THS in **A-B**. σ^70^V2TcR system & **C-D.** σ^70^V3-2LacI system. Time course of bioluminescence production by the **E.** pσ^70^V2TcR & **F.** pσ^70^V3-2LacI expression systems; each bar represents the bioluminescence measurement every 3 hours from time 0 to 18 hours (bars in each set, left to right, are 0h, 3h, 6h, 9h, 12h, 15h, and 18h). Difference in bioluminescence production between the ON and OFF states in **G.** pσ^70^V2TcR & **H.** pσ^70^V3-2LacI expression systems. Fold-change difference in the luminescence emission of the recombinant cells containing the **I.** pσ^70^V2TcR and **J.** pσ^70^V3-2LacI expression systems vs the wild type DH10B. The cultures were induced after 3 hours of growth with aTc or IPTG accordingly. For all samples, the bioluminescence mean was normalized by the cell density (OD_600_). Error bars are +/- S.D., n=3. Relative Luminescence Units (RLU).

These results demonstrate that the reporter luciferase provides a more sensitive assessment of the expression levels in the uninduced state compared to fluorescent proteins like *mCardinal*. Importantly, the translational regulation provided by the toehold switches efficiently amplifies the fold change and reduces the basal expression levels of the σ^70^ expression systems, thereby improving the tight regulation of the promoters while maintaining their strength (Fig 6G - J). However, total repression was not achieved by the adaptation of the toehold switch since the *σ^70^*V2TcR THS expression system showed minimal (Fig. 6E), and the *σ^70^*V3-2LacI THS pronounced (Fig. 6F) leakage, thus prohibiting total control of the target recombinant genes.

Total gene expression control is crucial for regulating enzymatic reactions involved in the biosynthesis of natural products, as demonstrated with the production of terpenes presented in this study. The synthetic expression systems evaluated in this study were evaluated in the pJH0204 plasmid which contains the pColE1 origin of replication [23]. This plasmid has the advantage to integrate in the chromosome of *P. putida* and replicate in *E. coli* and *V. natriegens*, therefore it can be evaluated in multiple chassis without further manipulation [23]. However, one potential disadvantage of this plasmid in *E. coli* is the copy number. The pColE1 origin of replication produces 25–30 copies per cell in *E. coli* [55]. Therefore, to reduce basal expression and achieve tighter control over gene expression, we transferred the synthetic expression systems to the low copy plasmid pSC101, which has approximately 3 copies per cell [64]. In this configuration, both *σ^70^*V2TcR expression systems (with and without the THS) achieved total repression of the lux operon in the uninduced state (Fig. 6E). Activation by aTc successfully induced bioluminescence production. However, the THS-version produced ∼2-fold less light, indicating that the translational control provided by the THS may be incompletely relieved in the induced state (Fig. 6G, I). A similar trend was observed with the *σ^70^*V2LacI and *σ^70^*V3-2LacI expression systems in pSC101 (Fig. 6F, H, J). Although total repression was not achieved, the translational control provided by the toehold significantly reduced the basal expression levels and amplified the dynamic range of the strong *σ^70^*V2Lac promoter (Fig. 6H, J). We conclude that the combination of translational regulation and reduced plasmid copy number significantly improved the problem of leaky transcription from our promoters in *E. coli*.

In *P. putida*, the pJH0204 plasmid integrates at one specific location in the chromosome [35]. In this configuration, using *mCardinal* as the reporter system, the *σ^70^*V2TcR showed no leakage, while the *σ^70^*V2LacI exhibited minimal red fluorescence in the uninduced state [23]. However, traces of lycopene and β-carotene were detected in the uninduced state of the *σ^70^*V2TcR (Fig. 4C, D), and only β-carotene was produced in the dual system (Fig. 4F, G), thus corroborating the leakage of *σ^70^*V2LacI in this host. Since the transcriptional control provided by the regulatory proteins TetR and/or LacI proved insufficient to fully repress the *σ^70^*synthetic promoters, even at one copy per cell, we decided to incorporate translation repression in *P. putida*. The luciferase reporter system confirmed the leakage of the *σ^70^* expression systems in this host (Fig. 7A-B, Fig. S5), and the addition of the THS amplified the dynamic range and reduced the luminescence emission in the uninduced state by ∼2-fold in both expression systems (Fig. 7C, D). These results demonstrate that the transcriptional control provided by the regulatory proteins can be enhanced by translational control via THS *σ^70^*V2TcR in *P. putida*. As in *E. coli*, these two layers of control are not sufficient to completely turn off the *σ^70^*V2LacI expression system in *P. putida*. Unlike in *E. coli*, plasmid copy number is not a parameter that could be further optimized in *P. putida*.

**Figure 7.**
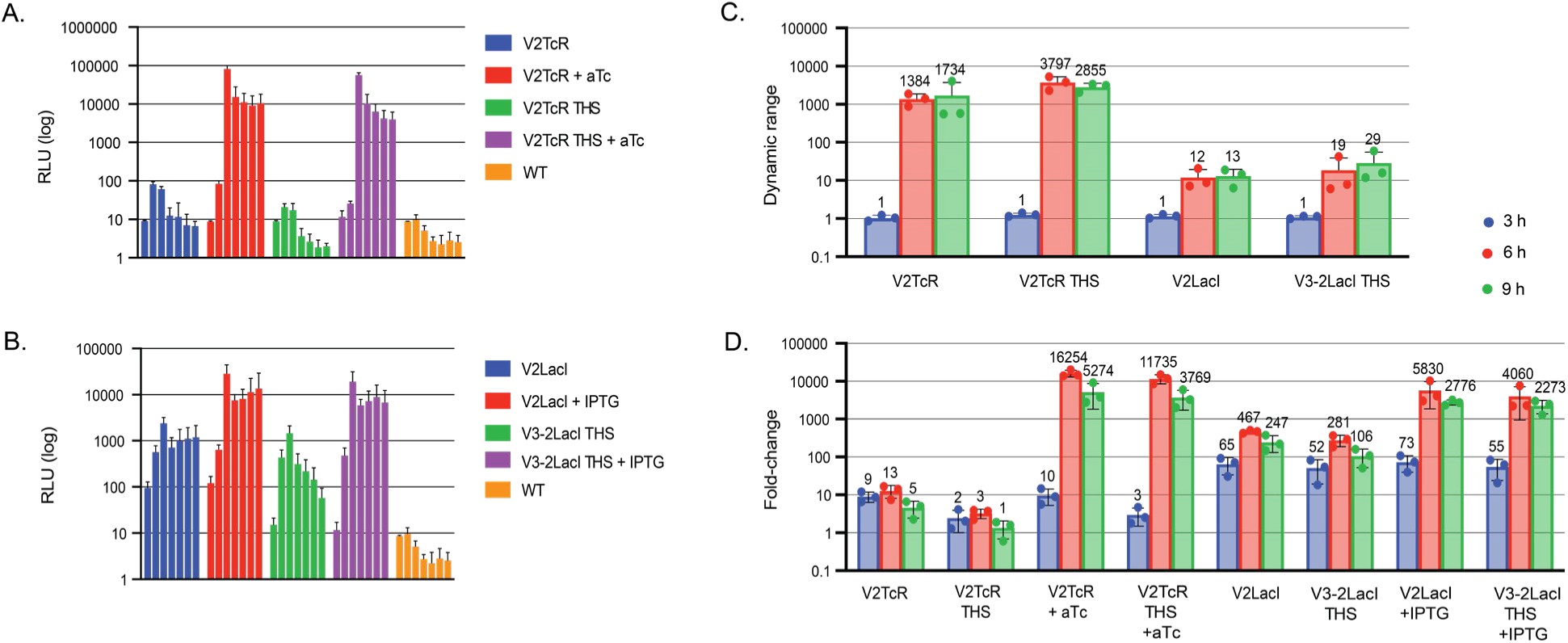
Comparison of the σ^70^ expression systems to the toehold versions controlling the expression of the lux operon in *P. putida*. Time course of bioluminescence production by the **A.** pσ^70^V2TcR and **B.** pσ^70^V3-2LacI expression systems; each bar represents the bioluminescence measurement every 3 hours from time 0 to 18 hours. Difference in bioluminescence production between the ON and OFF states in **C.** pσ^70^V2TcR and pσ^70^V3-2LacI expression systems. Fold-change difference in the light emission of the recombinant cells containing the **D**. pσ^70^V2TcR or and pσ^70^V3-2LacI expression systems vs the wild type *P. putida*. The cultures were induced after 3 hours of growth with aTc or IPTG accordingly. For all samples, the bioluminescence mean was normalized by the cell density (OD_600_). Error bars are +/- S.D., n=3. Relative Light Unites (RLU).

### Production of lycopene and β-carotene using the *σ^70^* THS control systems in *E. coli*

The data collected using the luciferase reporter system demonstrated that adapting the THS to the *σ^70^* expression systems effectively suppressed the basal expression of *σ^70^*V2TcR and significantly reduced the leakage of *σ^70^*V3-2LacI when tested in a low-copy plasmid (Fig. 6I, J). These findings enabled a re-evaluation of lycopene and β-carotene production, with the goal of independently controlling different steps within a multistep BGC. Previously, in *E. coli* expressing *crtEBI* and *crtEBIY* under the control of *σ^70^*V2TcR in a medium copy plasmid (pJH0204), the system essentially behaved as a constitutive promoter, with significant production of lycopene and β-carotene in the uninduced state, and less than 2-fold increase in product upon induction (Fig. 4C, D). To assess the ability of the THS designs to better regulate terpenoid production, lycopene and β-carotene production were evaluated separately using the *σ^70^*V2TcR-THS and *σ^70^*V3-2LacI-THS expression systems in the pSC101 vector (Fig. 8 A, B). With the *σ^70^*V2TcR-THS system, terpenoid production was completely abolished in the uninduced state. Upon the addition of aTc, lycopene levels reached titers of 553 µg/L, while β-carotene titers reached 455 µg/L (Fig. 8C, D). Thus, we were able to achieve complete OFF state control of these BCGs in *E. coli*.

**Figure 8.**
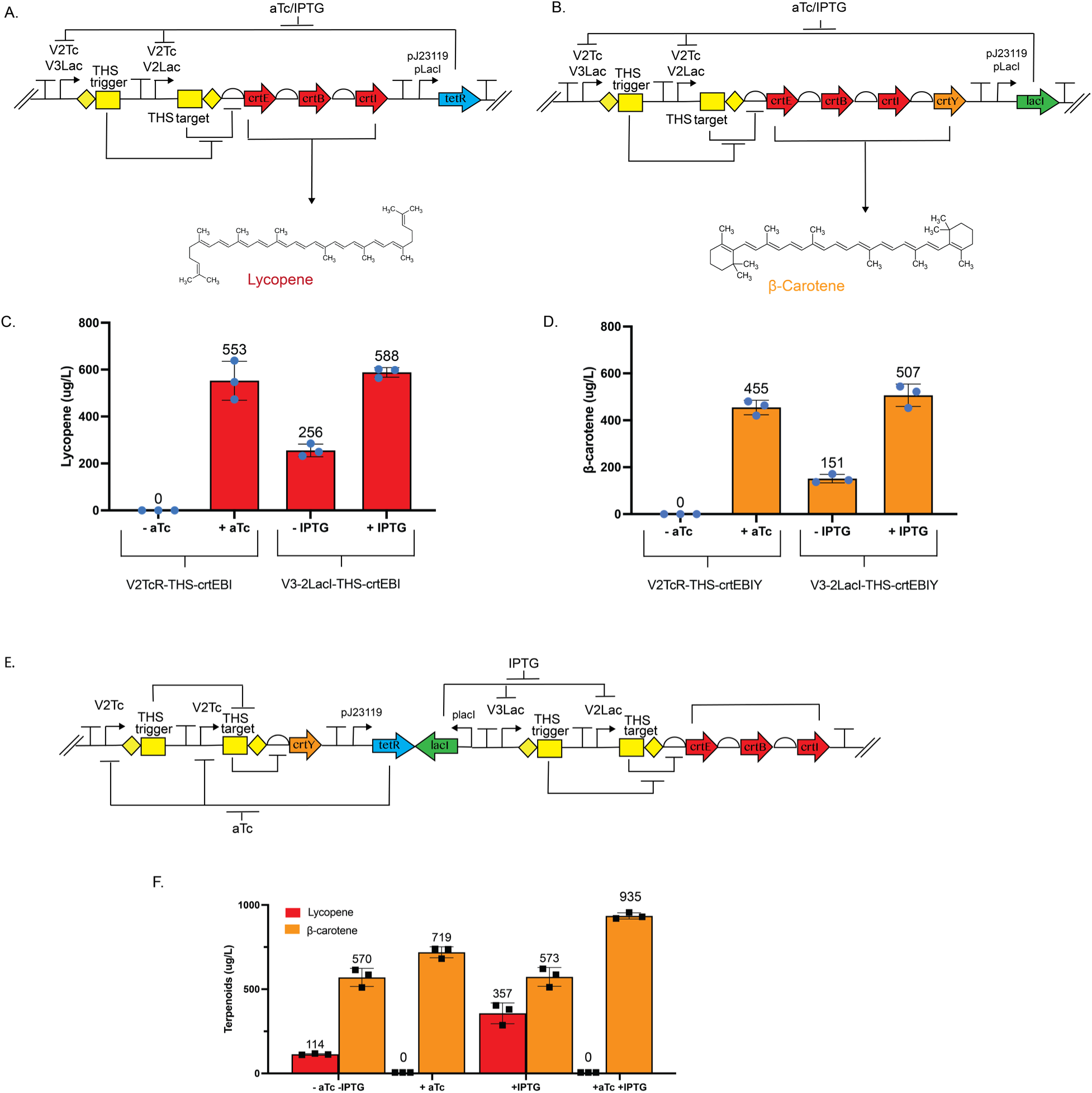
Production of lycopene and β-carotene in *E. coli* using the single and dual expression systems with THS transformed with pSC101 vector. Architecture of the σ^70^V2TcR σ^70^V3-2LacI-THS controlling the **A.** LYC operon and B. *crtEBIY* genes. Production of **C.** lycopene and **D.** β-carotene by the single expression systems. **E.** Gene circuit of the dual expression system with THSs **F.** Lycopene and β-carotene production by the dual expression system. Cultures were induced with aTc, IPTG or both inducers at OD_600_. 0.7 for 4 hours and terpenoids titers were measured by UHPLC. Error bars are +/- S.D., n=3.

The performance of the *σ^70^*V2/3LacI systems were not assessed in the pColE1 vector in our first set of experiments, due to its well documented leakage [23]. However, given the improvements made by the integration of the THS and measurements with the *lux* reporter, the *σ^70^*V3-2LacI-THS system was evaluated for terpenoid production. We still observed considerable levels of lycopene and β-carotene production in the uninduced state using *σ^70^*V3-2LacI-THS (Fig. 8C, D, and Fig. S6), though titers (256 µg/L for lycopene, and 151 µg/L for β-carotene) were approximately 1.8-fold lower than those observed in the uninduced state for *σ^70^*V2TcR in pColE1 (Fig. 4C, D). The addition of IPTG activated the terpenoid production reaching similar titers (588 µg/L for lycopene, and 507 µg/L for β-carotene) to the ones obtained with the *σ^70^*V2TcR-THS (Fig. 8C, D). This result represents a significant improvement in the performance of our *lac* promoter system in controlling a BGC.

The next step was to evaluate the capabilities of the dual expression system for producing both β-carotene and its intermediate, lycopene. In the earlier setup (Fig. 4E), the LYC operon (*crtEBI*) was controlled by *σ^70^*V2TcR, and *crtY* by *σ^70^*V3LacI. However, in this configuration, only β-carotene was produced due to significant leakage from both expression systems. This leakage prevented lycopene accumulation in the absence of *crtY* induction (Fig. 4F), as the high processivity of CrtY meant that the leaky expression was sufficient to produce enough enzyme to rapidly converted any lycopene produced into β-carotene (Fig.4G). To prevent a similar outcome in the new dual expression system, the *crtY* was placed under the control of the tight *σ^70^*V2TcR-THS expression system, and the LYC operon regulated by the more relaxed *σ^70^*V3-2LacI-THS expression system (Fig. 8E). The uninduced state of *σ^70^*V2TcR-THS-crtY / *σ^70^*V3-2LacI-THS-crtEBI revealed production of both terpenoids (Fig. 8F). We expected to see lycopene being produced, because we saw leaky lycopene from this promoter in Fig. 8C. However, we were surprised by the amount of β-carotene produced, as we expected *crtY* transcription and translation to be tightly controlled based on the results in Fig. 8C-D. Nonetheless, we observed a new behavior in this circuit that was not observed in the first iteration in Fig. 4G, that is, the accumulation of lycopene and β-carotene in the same cell (Fig. 8F). Similarly, when IPTG was added to induce *crtEBI*, the amount of lycopene accumulation increased. Addition of aTc to induce *crtY*, either alone or with IPTG, resulted in the rapid and complete conversion of all lycopene to β-carotene. The dual circuit seems to be producing carotenoids more efficiently, reaching a titer of 935 µg/L under dual induction conditions. The results suggest that leaky expression of *crtY* cannot be entirely eliminated in this configuration of our genetic circuit. However, the amount of CrtY enzyme produced has been decreased by the THS significantly so as to be incapable of metabolizing all available lycopene to β-carotene.

## CONCLUSION

The development of various bacterial chassis has garnered attention in pursuit of the goals of next-generation industrial biotechnology (NGIB) [65]. NGIB aims to utilize low-cost mixed substrates, minimize freshwater consumption, save energy, and implement long-lasting, continuous, intelligent processing methods [65]. Despite the numerous advantages of *E. coli*, such as its robust genetic toolbox and standardized cultivation parameters [66, 67], it falls short of achieving the goals of NGIB due to its sensitivity to harsh environments and limited metabolic capabilities [68, 69]. Alternative hosts may be more attractive to accomplish complex tasks. For example, the metabolic versatility of *P. putida* has demonstrated tolerance to organic solvents and stresses associated with industrial-scale cultivations, the ability to utilize lignin-derived compounds, sustain difficult redox reactions, and a relatively high GC-content similar to that of actinomycete-derived natural product BGCs [15, 70, 71]. The marine bacterium *V. natriegens*, inherently halophilic, offers interesting advantages as well, such as rapid growth and a versatile metabolism capable of utilizing a variety of substrates as carbon and energy sources [72–74]. Other non-model Gram-negative bacterial chassis also demonstrate the potential to outperform *E. coli* [75, 76]. However, each chassis necessitates the adaptation of distinct genetic tools. The *σ^70^* expression system showcased in this study, whether used individually or in combination, offers another tool to the ongoing challenge of developing numerous host organisms. It provides a versatile and robust gene expression platform with a broad host range. Consequently, proteins or BGCs of interest can be readily cloned into each *σ^70^* expression system and evaluated across various chassis, thereby expediting the optimization process.

In this study, we selected chassis bacteria commonly used by the scientific community and, through the straightforward incorporation of synthetic gene pathways for the biosynthesis of lycopene and β-carotene, achieved relatively high titers of these terpenoids using the *σ^70^*expression system. Notably, no media optimization was necessary to complete this task, highlighting a remarkable advantage offered by the *σ^70^* expression system. However, the production of terpenoids using the first generation of the *σ^70^* expression system revealed leakage in the OFF state, a feature imperceptible when using fluorescent proteins as a reporter system.

In this instance, the characterization of the regulatory systems with a multi-step enzymatic reporter (*lux*), was essential in our task to minimize leaky transcription. We acknowledge that differences in culture conditions between the *lux* study (performed in 96-well plates) and our terpene studies (grown in 5 mL cultures in 50 mL culture tubes) may account for some of the differences in performance between these experiments. Still, the general approach of employing an enzymatic reporter led to insights that improved our dual circuit performance when expressing a BGC. Approaches like decreasing plasmid copy number, weakening the RBS sequences, and introducing mutations in the consensus promoter sequences have demonstrated the ability to minimize leakage in the basal state, albeit at the cost of reducing the induced expression levels [77]. The strength and portability of the *σ^70^* expression systems reside in the specific nucleotide arrangement of the promoter and RBS DNA sequence [23]; therefore, altering these sequences undermines the value of the system. Instead, the adaptation of the riboswitch toehold, together with the use of a low-copy plasmid in *E. coli*, reduced the leakage of the *σ^70^*V2TcR THS to non-detectable levels and nearly abolished the basal expression levels of the *σ^70^*V3-2LacI. This approach not only amplified the dynamic range but also maintained the portability of the synthetic expression systems. This was demonstrated in *P. putida*, where the DNA was integrated into the chromosome, and both THS *σ^70^* expression systems maintained strong induction, with the *σ^70^*V2TcR THS reaching very low *lux* reporter signal. The second generation of the *σ^70^* expression system (with THS) remains digital, with ON and OFF state only, however, the emergence of plasmids like TULIP (TUnable Ligand Inducible Plasmid) with controllable copy number [64] could allow an analogue behavior of the versatile *σ^70^* expression systems.

While our dual inducible system remains imperfect, we did achieve the accumulation of a pathway intermediate (lycopene) using a combination of transcriptional and translational control. We were able to convert that intermediate to a final product (β-carotene) upon specific induction of the downstream enzyme (CrtY). Complementary engineering approaches, such as decreasing the processivity of CrtY, could be used to further tune the system and achieve complete and independent inducible control of intermediate substrates and products.

In conclusion, the expression system developed in this study expands the genetic toolkit of various promising Gram-negative chassis, thereby enhancing their capabilities to address fundamental biological questions and bolster their biotechnological applications. Ultimately, the *σ^70^*expression system has the potential to facilitate the expression of biosynthetic gene clusters (BGCs) that have previously been inaccessible or suboptimal due to the lack of appropriate gene tools or the limitations of model organisms such as *E. coli*.

## DECLARATIONS

### Ethics approval and consent to participate

Not applicable.

### Consent for publication

Not applicable.

### Availability of data and materials

The raw datasets used and/or analyzed during the current study are available from the corresponding author at reasonable request. All processed data generated or analyzed during this study are included in this published article and its supplementary information files.

### Competing interests

A.F.C.R and N.G.F have filed a U.S. patent application for the pσ^70^ *lac* and *tet* promoters described in this work.

### Funding

This work was supported by new faculty start-up funds from Worcester Polytechnic Institute to N.G.F. & the Sandooq Al Watan grant PRJ-SWARD-256 to A.G.

## Authors’ contributions

**Andres Felipe Carrillo Rincón:** Conceptualization (lead); data curation (lead); formal analysis (lead); investigation (lead); methodology (lead); resources (equal); validation (lead); visualization (lead); writing – original draft (lead); writing – review and editing (equal).

**Alexandra J. Cabral**: Methodology (supporting); writing – review and editing (supporting).

**Andras Gyorgy** Conceptualization (supporting); funding acquisition (lead); project administration (lead); resources (lead); supervision (supporting); writing – original draft (supporting); writing – review and editing (supporting).

**Natalie Gilks Farny:** Conceptualization (supporting); funding acquisition (lead); project administration (lead); resources (lead); supervision (lead); writing – original draft (equal); writing – review and editing (equal).

## Acknowledgements

We thank Adam Guss of Oak Ridge National Laboratory for *P. putida* strains and plasmids; Kevin Keating, Emma Tobin, and Eric Young of WPI for assistance with UHPLC experiments and for the gift of *crtEBIY* genes; Shimshon Belkin of Hebrew University of Jerusalem for the *luxCDABE* operon.

## Notes

### Competing Interest Statement

A.F.C.R and N.G.F have filed a U.S. patent application for the pσ70 lac and tet promoters described in this work. All other authors declare no competing interests.

### Summary of Updates

Text was revised in response to review, and new results were added, including Figure 8 and Supplementary figures S1-S6.

