## Supplementary Methods and Figures for "A dual-inducible control system for multistep biosynthetic pathways"

Detailed cloning strategy

The backbone of the pJH0204 vector was modified via Gibson to incorporate in the multicloning site the restriction sites BamHI, NdeI, HindIII, KpnI, EcoRI, PstI, XbaI, NcoI, XhoI and eliminate the EcoRI and NcoI sites from the kanamycin cassette, thus generating the plasmid pJH0204F. Then, the *σ^70^*V2TcR-*mCardinal* transcriptional unit was cloned into pJH0204F using BamHI and PstI. The new plasmid was digested with PstI and XbaI to accommodate the transcriptional unit placI-lacI amplified by PCR flanked by PstI and XbaI. Next, the plasmid was digested with NcoI and XhoI to accommodate *mCardinal* previously amplified and flanked with the same enzymes. Finally, the plasmid was opened with XbaI and NcoI and the V2Lac and V3Lac promoters were cloned via Gibson generating the plasmid p*σ^70^*V2TcR-*mCardinal*/V2LacI-*mCardinal* and p*σ^70^*V2TcR-*mCardinal*/V3LacI-*mCardinal* (Fig. 1). Next the *sfGFP* was cloned after the *σ^70^*V2Tc via NdeI and HindIII generating the plasmids p*σ^70^*V2TcR-*sfGFP*/V2LacI-*mCardinal* and p*σ^70^*V2TcR-*sfGFP*/V3LacI- *mCardinal*.

The *crtEBI* genes were PCR amplified adjusting an NdeI at the 5’ of crtE and a HindIII at the 3’ of the crtI and assembled in the pUC19 vector digested with BamHI and HindIII generating the plasmid pUC19-*crtEBI*. The *crtEBI* gene cluster was cloned into p*σ^70^*V2TcR via NdeI and HindIII producing the plasmid p*σ^70^* V2TcR-*crtEBI*. Further, *crtY* was cloned via Gibson into p*σ^70^*V2TcR-*crtEBI* digested with HindIII generating the plasmid p*σ^70^*V2TcR-*crtEBIY*. Finally, the *crtY* gene was PCR amplified flanked by NcoI and XhoI and cloned to generate the plasmids p*σ^70^*V2TcR-*crtEBI*/V2LacI-*crtY* and p*σ^70^*V2TcR-*crtEBI*/V3LacI-*crtY*. The DNA parts containing the *σ^70^*V2Tc-THS-trigger/V2Tc-THS-target and *σ^70^*V3Lac-THS-trigger/ V2Lac-THS-target were synthesized as gBlocks and incorporated via Gibson assembly into the plasmids pJH0204-V2TcR-*mCardinal* and pJH0204- *σ^70^*V3LacI-*mCardinal* previously digested with BamHI/NdeI and NdeI, respectively. Further, the p*σ^70^*V2TcR-THS-*mCardinal* and *σ^70^*V3-2LacI-THS-*mCardinal* plasmids were digested with NdeI/HindIII to replace the *mCardinal* with the *luxCDABE* operon, previously PCR amplified with appropriate overlapping tails and incorporated into the plasmid via Gibson assembly.

To produce the pSC101 plasmids the ampicillin cassette was replaced via Golden Gate assembly. Briefly, the pSC101 was PCR amplified in the outward direction with respect to the ampicillin cassette, creating the backbone amplicon, flanked with BsaI cut sites. The kanamycin cassette was PCR amplified flanked with BsaI cut sites that generate complementary sticky ends to the pSC101 backbone. Both DNA fragments were assembled using the standard Golden Gate protocol generating pSC101-Kn. The cassettes containing the *luxCDABE* operon with the different expression systems were digested with BamHI and XhoI and cloned into pSC101-Kn previously amplified with primers that incorporated the BamhI/XhoI restriction sites. The LYC and β-carotene operons were cloned in the p*σ^70^*V2TcR-THS and *σ^70^*V3-2LacI-THS vectors using NdeI and HindIII, and the DNA circuit containing the V2Tc-THS-crtEBI, V2Tc-THS-crtEBIY, V3-2Lac-THS-crtEBI and V3-2Lac-THS-crtEBIY were cloned to pSC101Kn via BamHI and XhoI. To produce the dual p*σ^70^* V2TcR-THS-*crtY* V3-2LacI-THS-*crtEBI* vector, the *crtY* was cloned in p*σ^70^*V2TcR-THS using NdeI and HindIII to later accommodate the V3-2LacI-THS-*crtEBI* circuit using XbaI and XhoI. The circuit V2TcR-THS-*crtY* V3-2LacI-THS-*crtEBI* was transferred to pSC101Kn using BamHI and XhoI.

DNA design of the toehold (trigger and target)

The toehold and trigger designs are adapted from reference [1] to our synthetic σ^70^ promoters.

**V3-2Lac toehold**

GGATCCGAGCTGTTGACACTTTATGCTTCCGGCTCGTATAATGTGTGTGGAATTGTGAGCGGATAACAAGTGGAATTGTGAGCGGATAACAATTTCACACAGGAAACAGAATCATgggactgactattctgtgcaatagtcagtaaagcagggataaacgagatagataagataagatagaggtttcagccaaaaaacttaagaccgccggtcttgtccactaccttgcagtaatgcggtggacaggatcggcggttttcttttctcttctcaaGAGCTGTTGACACTTTATGCTTCCGGCTCGTATAATGTGTGTGGAATTGTGAGCGGATAACAACGCAGTAAGAGAGGAATGTACATgggtcttatcttatctatctcgtttatccctgcatacagaaacagaggagatatgcaatgataaacgagaacctggcggcagcgcaaaagatgcgtCATatg

- V3Lac
- V2Lac
- Trigger
- T12 terminator
- Target

**V2Tc Toehold**

GGATCCGAGCTGTTGACAACTCTATCATTGATAGAGTTATAATGTTCCCTATCAGTGATAGAGACGCAGTAAGAGAGGAATGTACATgggactgactattctgtgcaatagtcagtaaagcagggataaacgagatagataagataagatagaggtttcagccaaaaaacttaagaccgccggtcttgtccactaccttgcagtaatgcggtggacaggatcggcggttttcttttctcttctcaaGAGCTGTTGACAACTCTATCATTGATAGAGTTATAATGTTCCCTATCAGTGATAGAGACGCAGTAAGAGAGGAATGTACATgggtcttatcttatctatctcgtttatccctgcatacagaaacagaggagatatgcaatgataaacgagaacctggcggcagcgcaaaagatgcgtCATat

- V2Tc
- Trigger
- T12 terminator
- Target

Table S1. Primers used in this study

| **Primer name** | **Sequence 5’ to 3’** | **Logic** |
| --- | --- | --- |
| 204F1_fwd | Ggcgcgtactccaaaaggatctaggtgaagatc | Add MCS to pJH204 (BamHI, NdeI, HindIII, KpnI, EcorI, PstI, XbaI, NcoI,XhoI) |
| 204F1_rev | gagttcttctgattagacaaaaaaaaggcgac |  |
| 204F2_fwd | Tttttttgtctaatcagaagaactcgtcaagaagg |  |
| 204F2_rev | gcccgacggcgaggatctcgtcgtgacgcatg |  |
| 204F3_fwd | cacgacgagatcctcgccgtcgggcatccgc |  |
| 204F3_rev | gtctagactgcaggaattcaagcttcatatgggatccggaccaaaacgAAA |  |
| 204F4_fwd | aagcttgaattcctgcagtctagaccatggctcgaggacgaacaataaggc |  |
| 204F4_rev | Cctagatccttttggagtacgcgcccgggga |  |
| Fw PstI-lacI | attgcCTGCAGTCACTGCCCGCTTTCCAGTCG | Add Psti/XbaI to placI-lacI |
| Rv XbaI-lacI | gcaatTCTAGAagtcaaaagcctccgaccggaggcttttgactCCCCGTGGCCGGGGGACTG |  |
| Fw cardi NcoI | gAATCCCATGGTGAGTAAGGGTGAGGAGCTCATTA | Add NcoI/XhoI to mCardinal |
| Rv cardi XhoI | TCAGTCTCGAGTTACTTGTCGTCATCATCCTTAT |  |
| XbaI-V3Lac-NcoI_fwd | gcctccggtcggaggcttttgacttCTAGAGAGCTGTTGACACTTTATG | Assemble V2/3LacI promoter to pJH204-V2TcR-mCardinal-lacI |
| XbaI-V3Lac-NcoI_rev | taatgagctcctcacccttactcacCATGGGATTCTGTTTCCTGTGTG |  |
| XbaI-V2Lac-NcoI_rev | taatgagctcctcacccttactcacCATGGGTACATTCCTCTCTTACTGC |  |
| Fw crtE | ccagtgaattcgagctcggtacccgggGATCCcatatgtatccgtttataaggac | Assemble LYC operon into pUC19 via Gibson |
| Rv crtE | ATCTACATTCCTCTCTTACTGCGTTAACTGACGGCAGCGA |  |
| Fw crtB | CGCAGTAAGAGAGGAATGTAGATATGAATAATCCGTCGTTACTCAATCATGCG |  |
| Rv crtB | TATATCTCCTTCTTAAAGTTAAACAAAATTATTTCAGAGCGGGCGCTGCCAG |  |
| Fw crtI | AATAATTTTGTTTAACTTTAAGAAGGAGATATAATGAAACCAACTACGGTAAT |  |
| Rv crtI | caggaaacagctatgaccatgattacgccaAGCTTcaaatcagatcctccag |  |
| FwYgibson | GTTTGATGCTGGAGGATCTGATTTGATCcAtCGCAGTAAGAGAGGAATGTAGATatggGAGCGGCTATGCAACCGC | Incorporate crtY into LYC operon via Gibson |
| RvYgibson | TCTTTTCTGGAATTTGGTACCGAGaagctTTCAACGATGAGTCGTCATAATGGCT |  |
| Fw crtY NcoI | AgttgcccatggGAGCGGCTATGCAACCGC | Add NcoI/XhoI to crtY |
| Rv crtY XhoI | AgttgcctcgagTCAACGATGAGTCGTCATAATGGCTT |  |
| FwluxToe | gcggcagcgcaaaagatgcgtcatatgactaaaaaaatttcattcattattaacggcca | Insert luxP into synthetic promoters via gibson |
| FwluxV2TcLac | CGCAGTAAGAGAGGAATGTAcatatgactaaaaaaatttcattcattattaacggcca |  |
| Rvluxter | CTGGAATTTGGTACCGAGAAGCTTtcaactatcaaacgcttcggttaagcttaaa |  |
| SJ202F | CCATTGGGTCTCGGCTTCCTATAAAAATAGGCGTATCACGAGGC | Incorporate BsaI sites to pSC101 |
| SJ202R | CCATTGGGTCTCGCTCCTGTTTGTTAGTCTTGATGCTTCACTG |  |
| Kan F Primer | CCATTGGGTCTCGGGAGTTGTGTCTCAAAATCTCTGATGTACATTG | Amplify kn cassette adding BsaI sites |
| Kan R Primer | CCATTGGGTCTCGAAGCGGAACGAAAAATCAATCTAAAGTATATATG |  |
| FwBamHI101 | ccgatGGATCCtactagagccaggcatcaaataaaa | Add BamHI and XhoI to pSC101 |
| RvXhoII101 | aacctCTCGAGattccagaaatcatccttagcgaaag |  |
| Fw crtY NdeI | cgataCATatggGAGCGGCTATGCAAC | add NdeI and HindIII tor crtY |
| Rv crtY HindIII | accttAAGCTTTCAACGATGAGTCGTCATAATGG |  |
| Fw Duet terminator XbaI | ccttttttcgttttggtccGTCTAGAGAGCT | add XbaI and XhoI V3-2Lac THS-crtEBI |
| Rv Duet terminator XhoI | TTTTCTGGAATTTGGTACCGAGAACTCGAGTCA |  |

DNA sequence of the LYC operon and crtY optimized for *E. coli* codon usage used in this study

crtE

ATGTATCCGTTTATAAGGACAGCCCGAATGACGGTCTGCGCAAAAAAACACGTTCATCTCACTCGCGATGCTGCGGAGCAGTTACTGGCTGATATTGATCGACGCCTTGATCAGTTATTGCCCGTGGAGGGAGAACGGGATGTTGTGGGTGCCGCGATGCGTGAAGGTGCGCTGGCACCGGGAAAACGTATTCGCCCCATGTTGCTGTTGCTGACCGCCCGCGATCTGGGTTGCGCTGTCAGCCATGACGGATTACTGGATTTGGCCTGTGCGGTGGAAATGGTCCACGCGGCTTCGCTGATCCTTGACGATATGCCCTGCATGGACGATGCGAAGCTGCGGCGCGGACGCCCTACCATTCATTCTCATTACGGAGAGCATGTGGCAATACTGGCGGCGGTTGCCTTGCTGAGTAAAGCCTTTGGCGTAATTGCCGATGCAGATGGCCTCACGCCGCTGGCAAAAAATCGGGCGGTTTCTGAACTGTCAAACGCCATCGGCATGCAAGGATTGGTTCAGGGTCAGTTCAAGGATCTGTCTGAAGGGGATAAGCCGCGCAGCGCTGAAGCTATTTTGATGACGAATCACTTTAAAACCAGCACGCTGTTTTGTGCCTCCATGCAGATGGCCTCGATTGTTGCGAATGCCTCCAGCGAAGCGCGTGATTGCCTGCATCGTTTTTCACTTGATCTTGGTCAGGCATTTCAACTGCTGGACGATTTGACCGATGGCATGACCGACACCGGTAAGGATAGCAATCAGGACGCCGGTAAATCGACGCTGGTCAATCTGTTAGGCCCGAGGGCGGTTGAAGAACGTCTGAGACAACATCTTCAGCTTGCCAGTGAGCATCTCTCTGCGGCCTGCCAACACGGGCACGCCACTCAACATTTTATTCAGGCCTGGTTTGACAAAAAACTCGCTGCCGTCAGTTAA

crtB

ATGAATAATCCGTCGTTACTCAATCATGCGGTCGAAACGATGGCAGTTGGCTCGAAAAGTTTTGCGACAGCCTCAAAGTTATTTGATGCAAAAACCCGGCGCAGCGTACTGATGCTCTACGCCTGGTGCCGCCATTGTGACGATGTTATTGACGATCAGACGCTGGGCTTTCAGGCCCGGCAGCCTGCCTTACAAACGCCCGAACAACGTCTGATGCAACTTGAGATGAAAACGCGCCAGGCCTATGCAGGATCGCAGATGCACGAACCGGCGTTTGCGGCTTTTCAGGAAGTGGCTATGGCTCATGATATCGCCCCGGCTTACGCGTTTGATCATCTGGAAGGCTTCGCCATGGATGTACGCGAAGCGCAATACAGCCAACTGGATGATACGCTGCGCTATTGCTATCACGTTGCAGGCGTTGTCGGCTTGATGATGGCGCAAATCATGGGCGTGCGGGATAACGCCACGCTGGACCGCGCCTGTGACCTTGGGCTGGCATTTCAGTTGACCAATATTGCTCGCGATATTGTGGACGATGCGCATGCGGGCCGCTGTTATCTGCCGGCAAGCTGGCTGGAGCATGAAGGTCTGAACAAAGAGAATTATGCGGCACCTGAAAACCGTCAGGCGCTGAGCCGTATCGCCCGTCGTTTGGTGCAGGAAGCAGAACCTTACTATTTGTCTGCCACAGCCGGCCTGGCAGGGTTGCCCCTGCGTTCCGCCTGGGCAATCGCTACGGCGAAGCAGGTTTACCGGAAAATAGGTGTCAAAGTTGAACAGGCCGGTCAGCAAGCCTGGGATCAGCGGCAGTCAACGACCACGCCCGAAAAATTAACGCTGCTGCTGGCCGCCTCTGGTCAGGCCCTTACTTCCCGGATGCGGGCTCATCCTCCCCGCCCTGCGCATCTCTGGCAGCGCCCGCTCTGA

crtI

ATGAAACCAACTACGGTAATTGGTGCAGGCTTCGGTGGCCTGGCACTGGCAATTCGTCTACAAGCTGCGGGGATTCCCGTCTTACTGCTTGAACAACGTGATAAACCCGGCGGTCGGGCTTATGTCTACGAGGATCAGGGGTTTACCTTTGATGCAGGCCCGACGGTTATCACCGATCCCAGTGCCATTGAAGAACTGTTTGCACTGGCAGGAAAACAGTTAAAAGAGTATGTCGAACTGCTGCCGGTTACGCCGTTTTACCGCCTGTGTTGGGAGTCAGGGAAGGTCTTTAATTACGATAACGATCAAACCCGGCTCGAAGCGCAGATTCAGCAGTTTAATCCCCGCGATGTCGAAGGTTATCGTCAGTTTCTGGACTATTCACGCGCGGTGTTTAAAGAAGGCTATCTAAAGCTCGGTACTGTCCCTTTTTTATCGTTCAGAGACATGCTTCGCGCCGCACCTCAACTGGCGAAACTGCAGGCATGGAGAAGCGTTTACAGTAAGGTTGCCAGTTACATCGAAGATGAACATCTGCGCCAGGCGTTTTCTTTCCACTCGCTGTTGGTGGGCGGCAATCCCTTCGCCACCTCATCCATTTATACGTTGATACACGCGCTGGAGCGTGAGTGGGGCGTCTGGTTTCCGCGTGGCGGCACCGGCGCATTAGTTCAGGGGATGATAAAGCTGTTTCAGGATCTGGGTGGCGAAGTCGTGTTAAACGCCAGAGTCAGCCACATGGAAACGACAGGAAACAAGATTGAAGCCGTGCATTTAGAGGACGGTCGCAGGTTCCTGACGCAAGCCGTCGCGTCAAATGCAGATGTGGTTCATACCTATCGCGACCTGTTAAGCCAGCACCCTGCCGCGGTTAAGCAGTCCAACAAACTGCAGACTAAGCGCATGAGTAACTCTCTGTTTGTGCTCTATTTTGGTTTGAATCACCATCATGATCAGCTCGCGCATCACACGGTTTGTTTCGGCCCGCGTTACCGCGAGCTGATTGACGAAATTTTTAATCATGATGGCCTCGCAGAGGACTTCTCACTTTATCTGCACGCGCCCTGTGTCACGGATTCGTCACTGGCGCCTGAAGGTTGCGGCAGTTACTATGTGTTGGCGCCGGTGCCGCATTTAGGCACCGCGAACCTCGACTGGACGGTTGAGGGGCCAAAACTACGCGACCGTATTTTTGCGTACCTTGAGCAGCATTACATGCCTGGCTTACGGAGTCAGCTGGTCACGCACCGGATGTTTACGCCGTTTGATTTTCGCGACCAGCTTAATGCCTATCATGGCTCAGCCTTTTCTGTGGAGCCCGTTCTTACCCAGAGCGCCTGGTTTCGGCCGCATAACCGCGATAAAACCATTACTAATCTCTACCTGGTCGGCGCAGGCACGCATCCCGGCGCAGGCATTCCTGGCGTCATCGGCTCGGCAAAAGCGACAGCAGGTTTGATGCTGGAGGATCTGATTTGA

crtY

atggGAGCGGCTATGCAACCGCATTATGATCTGATTCTCGTGGGGGCTGGACTCGCGAATGGCCTTATCGCCCTGCGTCTCCAGCAGCAGCAACCTGATATGCGTATTTTGCTTATCGACGCCGCACCCCAGGCGGGCGGGAATCATACGTGGTCATTTCACCACGATGATTTGACTGAGAGCCAACATCGTTGGATAGCTCCGCTGGTGGTTCATCACTGGCCCGACTATCAGGTACGCTTTCCCACACGCCGTCGTAAGCTGAACAGCGGCTACTTTTGTATTACTTCTCAGCGTTTCGCTGAGGTTTTACAGCGACAGTTTGGCCCGCACTTGTGGATGGATACCGCGGTCGCAGAGGTTAATGCGGAATCTGTTCGGTTGAAAAAGGGTCAGGTTATCGGTGCCCGCGCGGTGATTGACGGGCGGGGTTATGCGGCAAATTCAGCACTGAGCGTGGGCTTCCAGGCGTTTATTGGCCAGGAATGGCGATTGAGCCACCCGCATGGTTTATCGTCTCCCATTATCATGGATGCCACGGTCGATCAGCAAAATGGTTATCGCTTCGTGTACAGCCTGCCGCTCTCGCCGACCAGATTGTTAATTGAAGATACGCACTATATTGATAATGCGACATTAGATCCTGAATGCGCGCGGCAAAATATTTGCGACTATGCCGCGCAACAGGGTTGGCAGCTTCAGACACTGCTGCGAGAAGAACAGGGCGCCTTACCCATTACTCTGTCGGGCAATGCCGACGCATTCTGGCAGCAGCGCCCCCTGGCCTGTAGTGGATTACGTGCCGGTCTGTTCCATCCTACCACCGGCTATTCACTGCCGCTGGCGGTTGCCGTGGCCGACCGCCTGAGTGCACTTGATGTCTTTACGTCGGCCTCAATTCACCATGCCATTACGCATTTTGCCCGCGAGCGCTGGCAGCAGCAGGGCTTTTTCCGCATGCTGAATCGCATGCTGTTTTTAGCCGGACCCGCCGATTCACGCTGGCGGGTTATGCAGCGTTTTTATGGTTTACCTGAAGATTTAATTGCCCGTTTTTATGCGGGAAAACTCACGCTGACCGATCGGCTACGTATTCTGAGCGGCAAGCCGCCTGTTCCGGTATTAGCAGCATTGCAAGCCATTATGACGACTCATCGTTGA

DNA sequence of the lux operon used in this study (from ref. [2])

luxC

atgactaaaaaaatttcattcattattaacggccaggttgaaatctttcccgaaagtgatgatttagtgcaatccattaattttggtgataatagtgtttacctgccaatattgaatgactctcatgtaaaaaacattattgattgtaatggaaataacgaattacggttgcataacattgtcaattttctctatacggtagggcaaagatggaaaaatgaagaatactcaagacgcaggacatacattcgtgacttaaaaaaatatatgggatattcagaagaaatggctaagctagaggccaattggatatctatgattttatgttctaaaggcggcctttatgatgttgtagaaaatgaacttggttctcgccatatcatggatgaatggctacctcaggatgaaagttatgttcgggcttttccgaaaggtaaatctgtacatctgttggcaggtaatgttccattatctgggatcatgtctatattacgcgcaattttaactaagaatcagtgtattataaaaacatcgtcaaccgatccttttaccgctaatgcattagcgttaagttttattgatgtagaccctaatcatccgataacgcgctctttatctgttatatattggccccaccaaggtgatacatcactcgcaaaagaaattatgcgacatgcggatgttattgtcgcttggggagggccagatgcgattaattgggcggtagagcatgcgccatcttatgctgatgtgattaaatttggttctaaaaagagtctttgcattatcgataatcctgttgatttgacgtccgcagcgacaggtgcggctcatgatgtttgtttttacgatcagcgagcttgtttttctgcccaaaacatatattacatgggaaatcattatgaggaatttaagttagcgttgatagaaaaacttaatctatatgcgcatatattaccgaatgccaaaaaagattttgatgaaaaggcggcctattctttagttcaaaaagaaagcttgtttgctggattaaaagtagaggtggatattcatcaacgttggatgattattgagtcaaatgcaggtgtggaatttaatcaaccacttggcagatgtgtgtaccttcatcacgtcgataatattgagcaaatattgccttatgttcaaaaaaataagacgcaaaccatatctatttttccttgggagtcatcatttaaatatcgagatgcgttagcattaaaaggtgcggaaaggattgtagaagcaggaatgaataacatatttcgagttggtggatctcatgacggaatgagaccgttgcaacgattagtgacatatatttctcatgaaaggccatctaactatacggctaaggatgttgcggttgaaatagaacagactcgattcctggaagaagataagttccttgtatttgtcccataa

luxD

atggaaaatgaatcaaaatataaaaccatcgaccacgttatttgtgttgaaggaaataaaaaaattcatgtttgggaaacgctgccagaagaaaacagcccaaagagaaagaatgccattattattgcgtctggttttgcccgcaggatggatcattttgctggtctggcggaatatttatcgcggaatggatttcatgtgatccgctatgattcgcttcaccacgttggattgagttcagggacaattgatgaatttacaatgtctataggaaagcagagcttgttagcagtggttgattggttaactacacgaaaaataaataacttcggtatgttggcttcaagcttatctgcgcggatagcttatgcaagcctatctgaaatcaatgcttcgtttttaatcaccgcagtcggtgttgttaacttaagatattctcttgaaagagctttagggtttgattatctcagtctacccattaatgaattgccggataatctagattttgaaggccataaattgggtgctgaagtctttgcgagagattgtcttgattttggttgggaagatttagcttctacaattaataacatgatgtatcttgatataccgtttattgcttttactgcaaataacgataattgggtcaagcaagatgaagttatcacattgttatcaaatattcgtagtaatcgatgcaagatatattctttgttaggaagttcgcatgacttgagtgaaaatttagtggtcctgcgcaatttttatcaatcggttacgaaagccgctatcgcgatggataatgatcatctggatattgatgttgatattactgaaccgtcatttgaacatttaactattgcgacagtcaatgaacgccgaatgagaattgagattgaaaatcaagcaatttctctgtcttaa

luxA

atgaaatttggaaactttttgcttacataccaacctccccaattttctcaaacagaggtaatgaaacgtttggttaaattaggtcgcatctctgaggagtgtggttttgataccgtatggttactggagcatcatttcacggagtttggtttgcttggtaacccttatgtcgctgctgcatatttacttggcgcgactaaaaaattgaatgtaggaactgccgctattgttcttcccacagcccatccagtacgccaacttgaagatgtgaatttattggatcaaatgtcaaaaggacgatttcggtttggtatttgccgagggctttacaacaaggactttcgcgtattcggcacagatatgaataacagtcgcgccttagcggaatgctggtacgggctgataaagaatggcatgacagagggatatatggaagctgataatgaacatatcaagttccataaggtaaaagtaaaccccgcggcgtatagcagaggtggcgcaccggtttatgtggtggctgaatcagcttcgacgactgagtgggctgctcaatttggcctaccgatgatattaagttggattataaatactaacgaaaagaaagcacaacttgagctttataatgaagtggctcaagaatatgggcacgatattcataatatcgaccattgcttatcatatataacatctgtagatcatgactcaattaaagcgaaagagatttgccggaaatttctggggcattggtatgattcttatgtgaatgctacgactatttttgatgattcagaccaaacaagaggttatgatttcaataaagggcagtggcgtgactttgtattaaaaggacataaagatactaatcgccgtattgattacagttacgaaatcaatcccgtgggaacgccgcaggaatgtattgacataattcaaaaagacattgatgctacaggaatatcaaatatttgttgtggatttgaagctaatggaacagtagacgaaattattgcttccatgaagctcttccagtctgatgtcatgccatttcttaaagaaaaacaacgttcgctattatattag

luxB

atgaaatttggattgttcttccttaacttcatcaattcaacaactgttcaagaacaaagtatagttcgcatgcaggaaataacggagtatgttgataagttgaattttgaacagattttagtgtatgaaaatcatttttcagataatggtgttgtcggcgctcctctgactgtttctggttttctgctcggtttaacagagaaaattaaaattggttcattaaatcacatcattacaactcatcatcctgtcgccatagcggaggaagcttgcttattggatcagttaagtgaagggagatttattttagggtttagtgattgcgaaaaaaaagatgaaatgcatttttttaatcgcccggttgaatatcaacagcaactatttgaagagtgttatgaaatcattaacgatgctttaacaacaggctattgtaatccagataacgatttttatagcttccctaaaatatctgtaaatccccatgcttatacgccaggcggacctcggaaatatgtaacagcaaccagtcatcatattgttgagtgggcggccaaaaaaggtattcctctcatctttaagtgggatgattctaatgatgttagatatgaatatgctgaaagatataaagccgttgcggataaatatgacgttgacctatcagagatagaccatcagttaatgatattagttaactataacgaagatagtaataaagctaaacaagagacgcgtgcatttattagtgattatgttcttgaaatgcaccctaatgaaaatttcgaaaataaacttgaagaaataattgcagaaaacgctgtcggaaattatacggagtgtataactgcggctaagttggcaattgaaaagtgtggtgcgaaaagtgtattgctgtcctttgaaccaatgaatgatttgatgagccaaaaaaatgtaatcaatattgttgatgataatattaagaagtaccacatggaatatacctaa

luxE

atgaagggtataaaagagtatgacagcagtgctgccatactttctaatattatcttgaggagtaaaacaggtatgacttcatacgttgataaacaagaaattacagcaagctcagaaattgatgatttgattttttcgagcgatccattagtgtggtcttacgacgagcaggaaaaaatcagaaagaaacttgtgcttgatgcatttcgtaatcattataaacattgtcgagaatatcgtcactactgtcaggcacacaaagtagatgacaatattacggaaattgatgacatacctgtattcccaacatcggtttttaagtttactcgcttattaacttctcaggaaaacgagattgaaagttggtttaccagtagcggcacgaatggtttaaaaagtcaggtggcgcgtgacagattaagtattgagagactcttaggctctgtgagttatggcatgaaatatgttggtagttggtttgatcatcaaatagaattagtcaatttgggaccagatagatttaatgctcataatatttggtttaaatatgttatgagtttggtggaattgttatatcctacgacatttaccgtaacagaagaacgaatagattttgttaaaacattgaatagtcttgaacgaataaaaaatcaagggaaagatctttgtcttattggttcgccatactttatttatttactctgccattatatgaaagataaaaaaatctcattttctggagataaaagcctttatatcataaccggaggcggctggaaaagttacgaaaaagaatctctgaaacgtgatgatttcaatcatcttttatttgatactttcaatctcagtgatattagtcagatccgagatatatttaatcaagttgaactcaacacttgtttctttgaggatgaaatgcagcgtaaacatgttccgccgtgggtatatgcgcgagcgcttgatcctgaaacgttgaaacctgtacctgatggaacgccggggttgatgagttatatggatgcgtcagcaaccagttatccagcatttattgttaccgatgatgtcgggataattagcagagaatatggtaagtatcccggcgtgctcgttgaaattttacgtcgcgtcaatacgaggacgcagaaagggtgtgctttaagcttaaccgaagcgtttgatagttga

**Supplementary Figures**

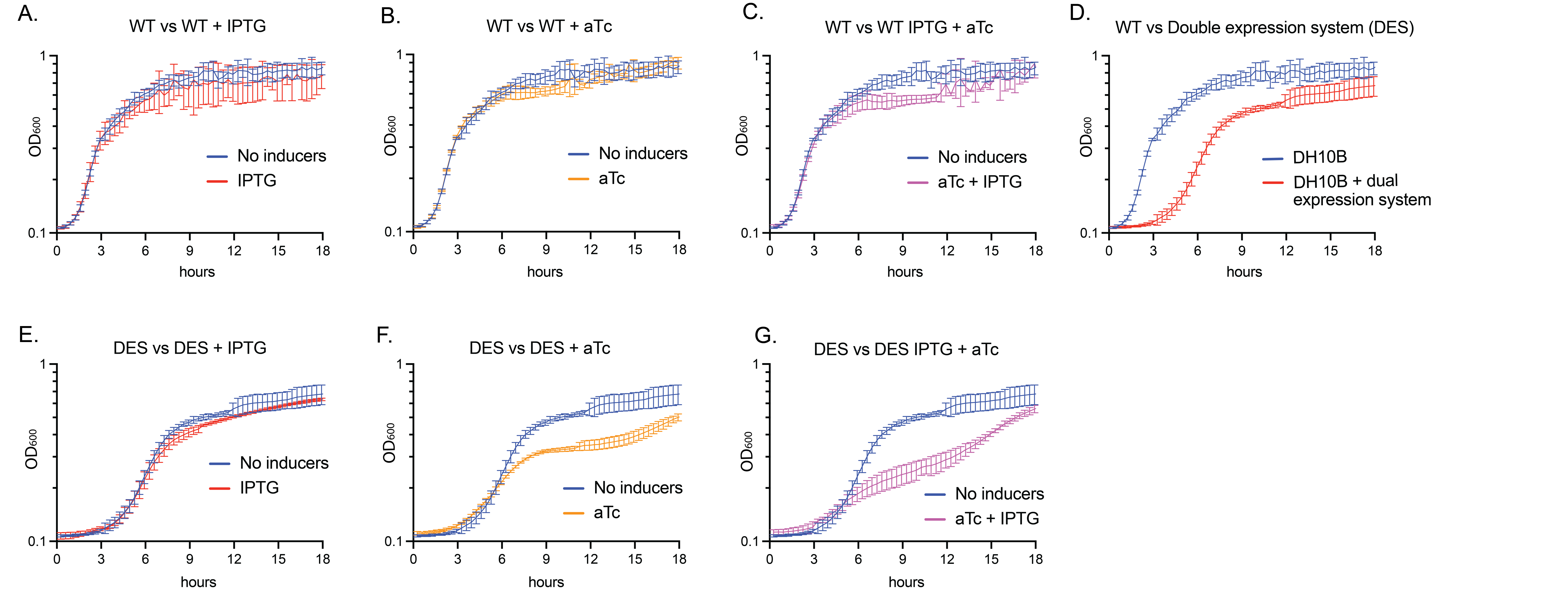

**Supplementary Figure S1.** Comparison of the growth rate dynamics of the wild type strain *E.coli* DH10B cultured in LB media exposed to **A.** IPTG **B.** aTc **C.** IPTG + aTc & **D.** the recombinant strain containing the *mCardinal* double expression system (DES: p*σ^70^* V2TcR-mCardinal/V3LacI-mCardinal version). Comparison of the growth rate dynamics of DES cultured in LB media exposed to **E.** IPTG **F.** aTc & **G.** IPTG + aTc. Error bars are +/- S.D., n=3

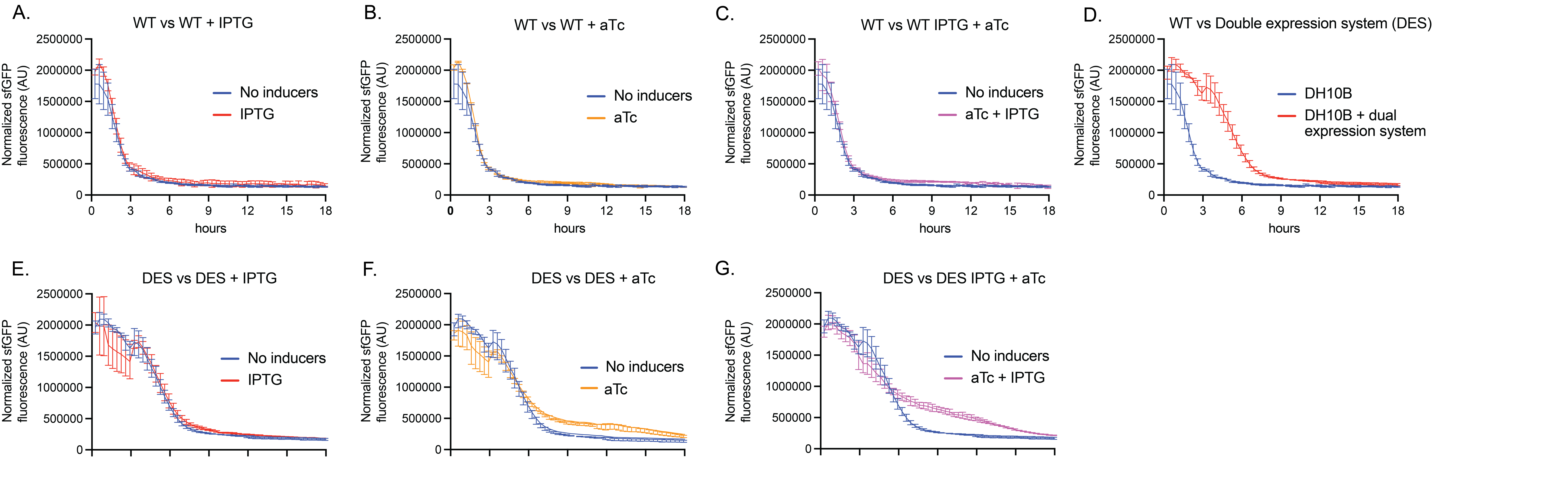

**Supplementary Figure S2.** Comparison of the green fluorescent emission normalized to OD_600_ of the wild type strain *E. coli* DH10B cultured in LB media exposed to **A.** IPTG **B.** aTc **C.** IPTG + aTc & **D.** the recombinant strain containing the mCardinal double expression system DES (p*σ^70^* V2TcR-mCardinal/V3LacI-mCardinal). Comparison of the green fluorescent emission normalized to OD_600_ of DES cultured in LB media exposed to **E.** IPTG **F.** aTc & **G.** IPTG + aTc. Error bars are +/- S.D., n=3

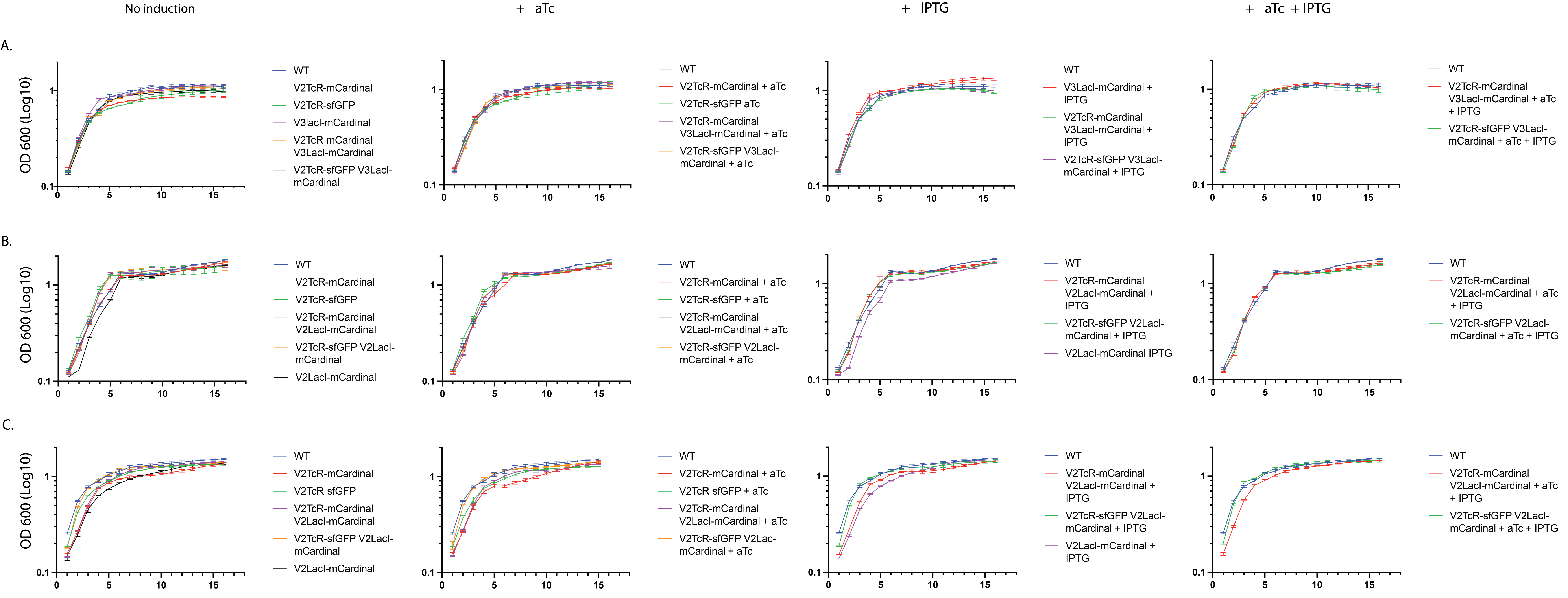

**Supplementary Figure S3.** Growth curves of the recombinant strains containing the single and double expression systems controlling the expression of sfGFP and/or mCardinal. A) E. coli B) P. putida & C) V. natriegens. Error bars are +/- S.D., n=3

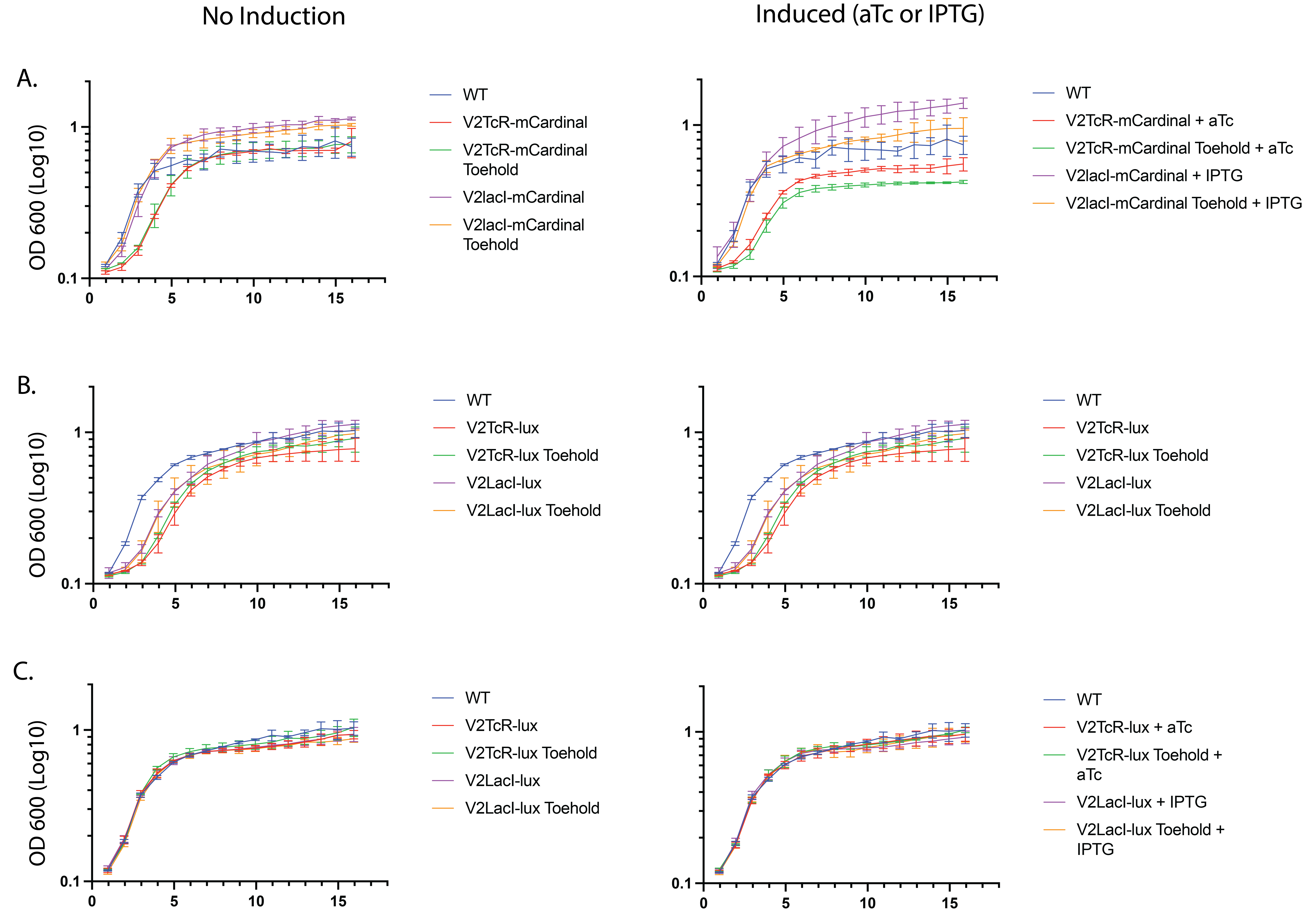

**Supplementary Figure S4.** Growth curves of the recombinant strains containing the single expression systems controlling the expression of mCardinal or the lux operon with or without the Toehold switch. A) *E. coli* with mCardinal in pColE1 ori B) *E. coli* with the lux operon in pColE1 ori & C) *E. coli* with the lux operon in pSC101 ori. Error bars are +/- S.D., n=3

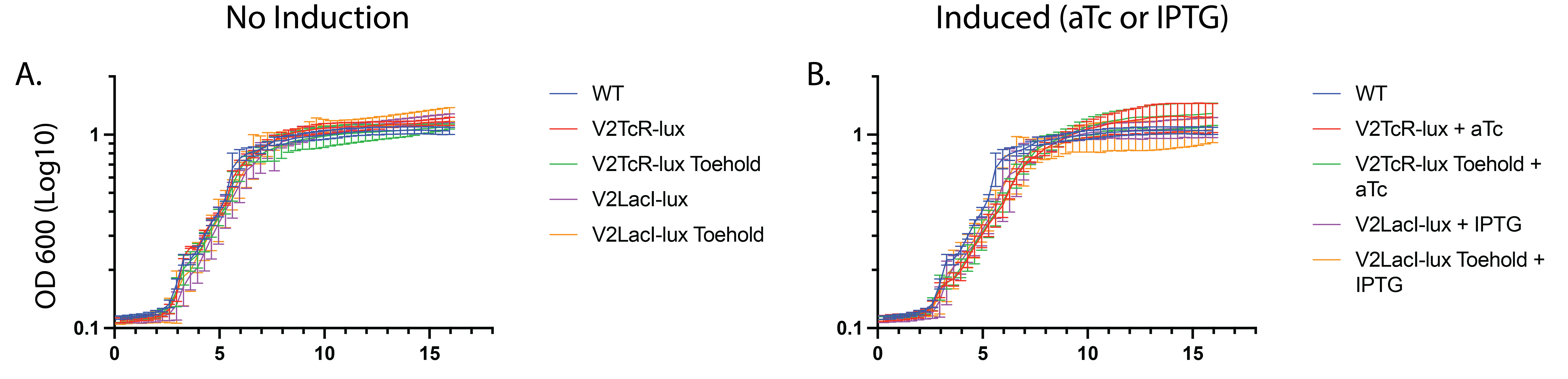

**Supplementary Figure S5.** Growth curves of the *P. putida* recombinant strains containing the single expression systems controlling the expression of the lux operon with or without the Toehold switch. A) Without the inducer B) With the inducers aTc or IPTG. Error bars are +/- S.D., n=3

A. pSC101Kn σ70 V2TcR-THS-crtY / V3-2LacI-THS-crtEBI

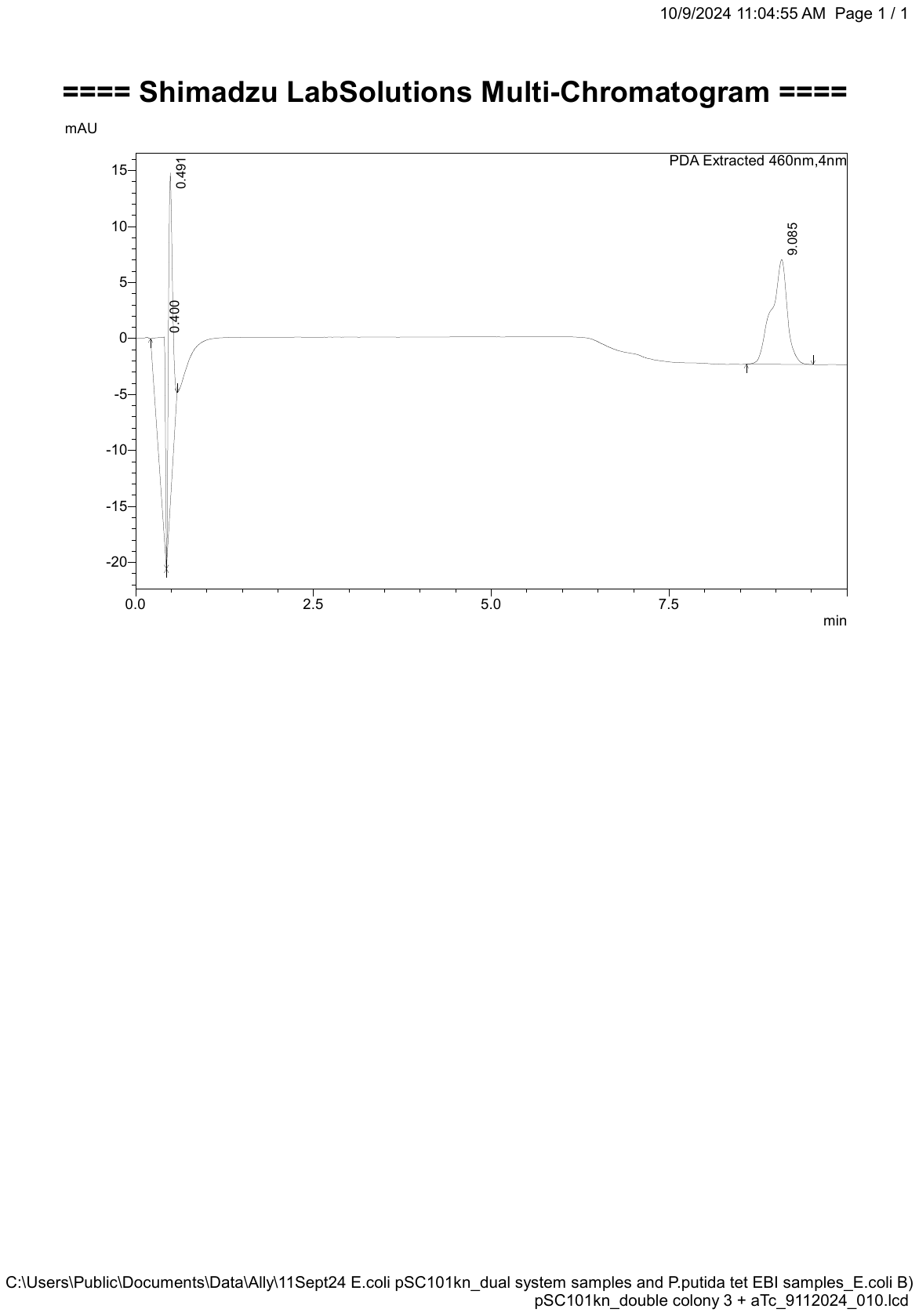

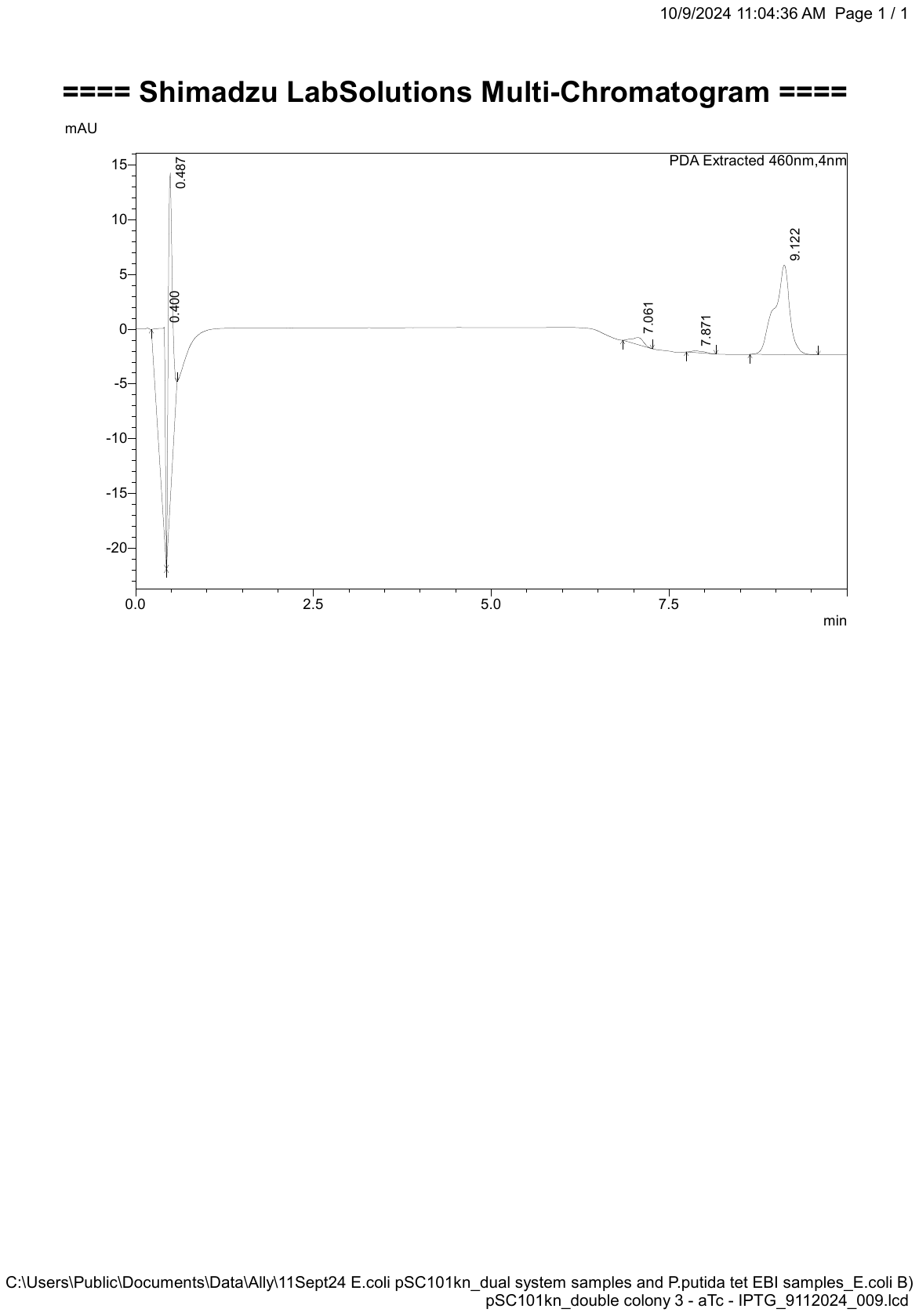

+aTc

-aTc -IPTG

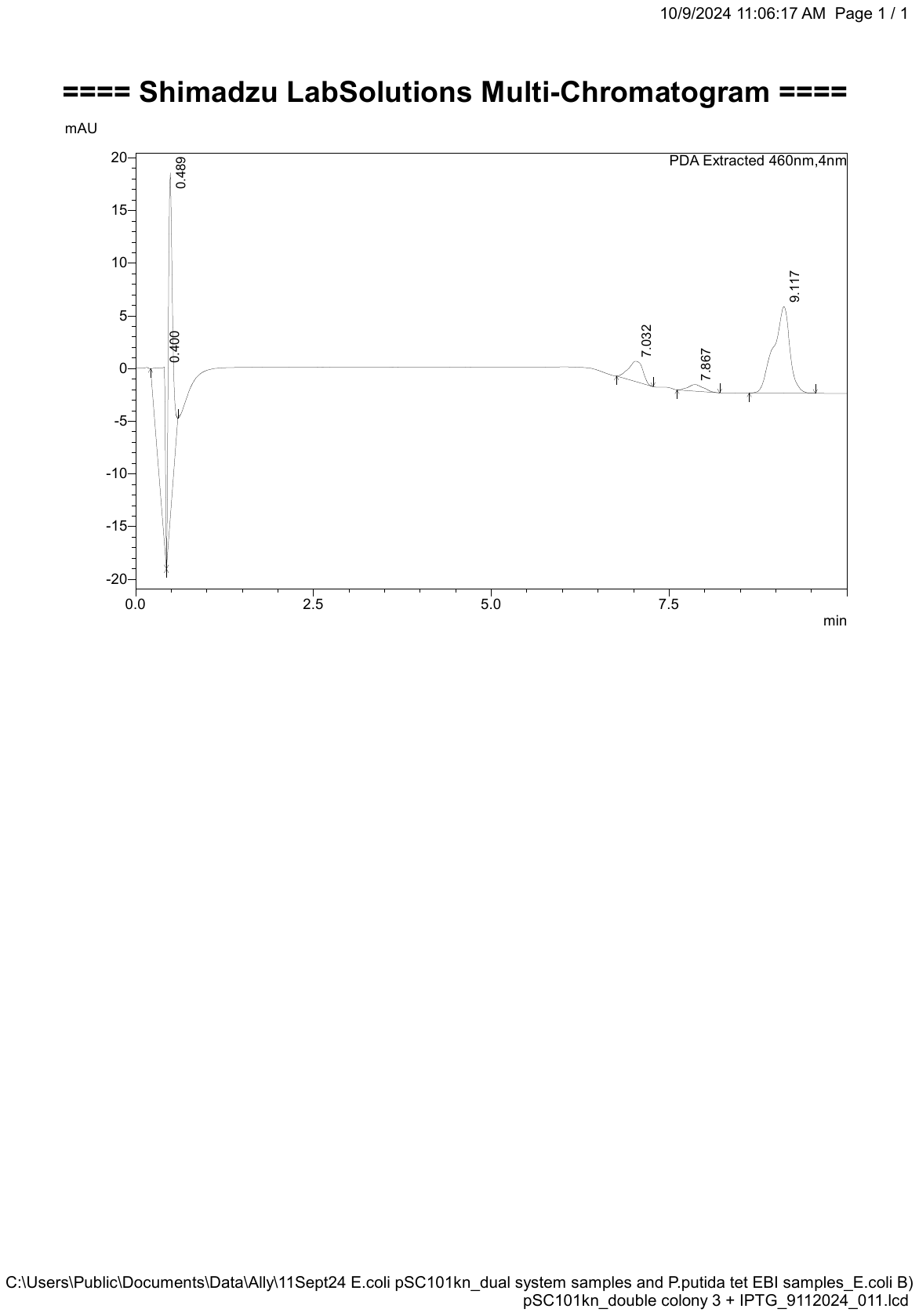

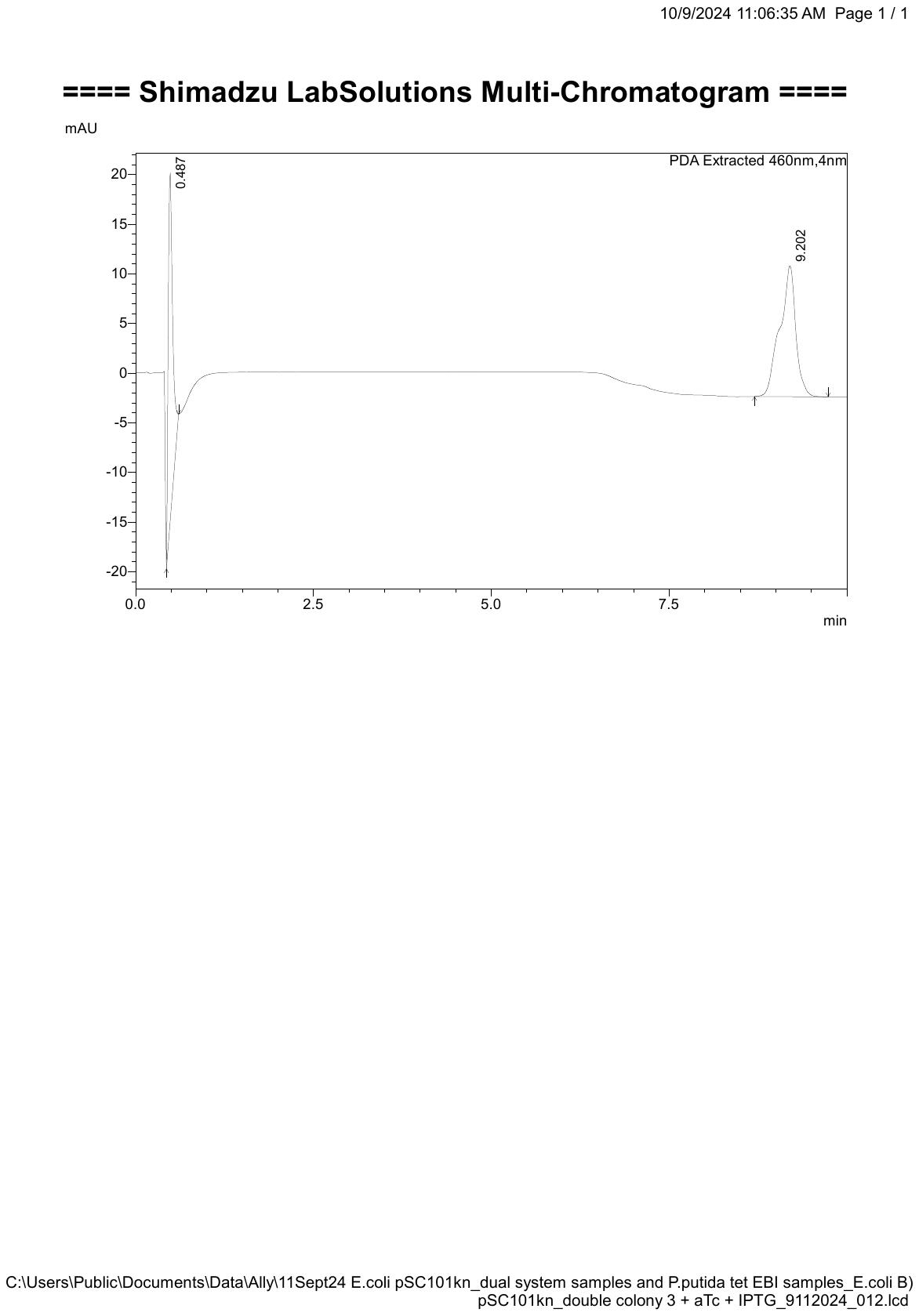

+aTc +IPTG

+IPTG

B.

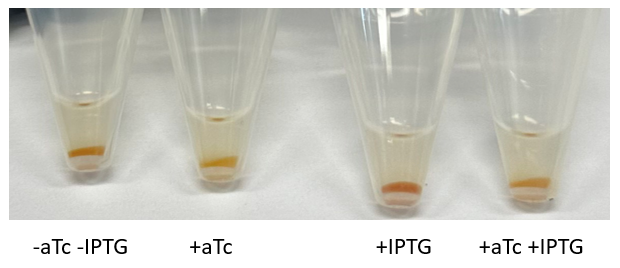

**Supplementary Figure S6.** Measurement of lycopene and β-carotene from the toehold-controlled dual inducible system (pSC101Kn σ70 V2TcR-THS-crtY V3-2LacI-Toehold-crtEBI) by HLPC. **A.** Representative chromatograms of HPLC samples measured in Figure 8. **B.** Representative cell pellets from measured samples. Darker red color is indicative of lycopene, whereas orange is β-carotene.
